# Extensive Copy Number Variation Explains Genome Size Variation in the Unicellular Zygnematophycean Alga, *Closterium peracerosum-strigosum-littorale* Complex

**DOI:** 10.1101/2023.05.13.540557

**Authors:** Yawako W. Kawaguchi, Yuki Tsuchikane, Keisuke Tanaka, Teruaki Taji, Yutaka Suzuki, Atsushi Toyoda, Motomi Ito, Yasuyuki Watano, Tomoaki Nishiyama, Hiroyuki Sekimoto, Takashi Tsuchimatsu

## Abstract

Genome sizes are known to vary within and among closely related species, but the knowledge about genomic factors contributing to the variation and their impacts on gene functions is limited to only a small number of species. This study identified a more than twofold heritable genome size variation among the unicellular Zygnematophycean alga, *Closterium peracerosum-strigosum-littorale* (*C. psl.*) complex, based on short-read sequencing analysis of 22 natural strains and F_1_ segregation analysis. Six *de novo* assembled genomes revealed that genome size variation is largely attributable to genome-wide copy number variation (CNV) among strains rather than mating type-linked genomic regions or specific repeat sequences such as rDNA. Notably, about 30% of genes showed CNV even between strains that can mate with each other. Transcriptome and gene ontology analysis demonstrated that CNV is distributed nonrandomly in terms of gene functions, such that CNV was more often observed in the gene set with stage-specific expression. Furthermore, in about 30% of these genes with CNV, the expression level does not increase proportionally with the gene copy number, suggesting presence of dosage compensation, which was overrepresented in genes involved in basic biological functions, such as translation. Nonrandom patterns in gene duplications and corresponding expression changes in terms of gene functions may contribute to maintaining the high level of CNV associated with extensive genome size variation in the *C. psl*. complex, despite its possible detrimental effects.

## Introduction

The genomes of eukaryotes vary tremendously in size, with more than a 200,000-fold variation among species (Gregory 2005). Genome size variation is mainly because of the different abundance of various repeat sequences, such as satellite DNA, transposable elements, and ribosomal genes (Kidwell 2002; Biémont 2008; Ågren and Wright 2011), while other factors, including heteromorphic sex chromosomes (Dolezel and Göhde 1995; Canon et al. 2000; Hjelmen et al. 2019), polyploidy, and aneuploidy (Rosato et al. 1998; Zhang et al. 2012; Poulíčková et al. 2014), have also been reported.

Substantial genome size variation within and among closely related species is relatively well documented in plants and animals (Šmarda et al. 2010; Long et al. 2013; Huang et al. 2014; Sun et al. 2018). In a few study systems, the genomic factors contributing to the variation, the inheritance pattern, and potential involvement of selection have been well characterized. In *Arabidopsis thaliana*, genome size ranges from 161 to 184 Mb among natural strains, showing an over 10% difference, which is largely explained by copy number variation (CNV) of 45S rDNA (Long et al. 2013). In the *Drosophila melanogaster* Genetic Reference Panel, genome size ranges from 169.7 to 192.8 Mb, and its skewed distribution toward the accumulation of large genomes suggests a greater constraint on genome reduction than expansion (Huang et al. 2014). Huang et al. (2014) also revealed that genome size is associated with polymorphic inversions. The rotifer *Brachionus asplanchnoidis* displays almost a twofold variation within a geographic population (Stelzer et al. 2019), and several large contiguous copy number variable regions, mainly composed of tandemly repeated satellite DNA, are associated with genome size (Stelzer et al. 2021). However, despite the prevalence of genome size variation within closely related species, the knowledge about the basic mechanisms and the inheritance pattern is still limited to only a few species.

In addition, how genome size variation would generally influence gene functions and phenotypes remains largely unknown. If genome size variation is partly due to CNV of genes, such variation altering gene dosage could potentially impact the expression, gene functions, and phenotypes, as gene dosage changes are often associated with detrimental effects such as disease, slow growth, and sensitivity to stress (Torres et al. 2007; Conrad et al. 2010; Cooper et al. 2011; Dephoure et al. 2014; Rice and McLysaght 2017). The prevalence of extensive genome size variation within species and among closely related species implies the presence of compensating mechanisms that buffer gene dosage changes and mitigate the imbalance. Recent studies detected dosage compensation in response to CNV among natural or artificial strains in yeast, plants, and humans (Huettel et al. 2008; Woodwark and Bateman 2011; Hose et al. 2015; Gasch et al. 2016; Zhang et al. 2017), but how generally dosage compensation is observed remains controversial (Torres et al. 2016), and so far most studies have focused on a limited number of species, in particular model organisms. We note that while the widely recognized dosage compensation of sex chromosomes in animals is mainly regulated epigenetically at the chromosome scale (Park and Koroda 2001; Distech 2012), gene-wise modulations of expression are also observed, which are called dosage compensation in this study as well (Guo and Birchler 1994; Hose et al. 2015; Gasch et al. 2016).

Here, we investigated the genome size variation, its characterization at the genome sequence level, the pattern of inheritance, and the functional consequences of the variation in a group of unicellular isogamous cosmopolitan algae, the *Closterium peracerosum-strigosum-littorale* (*C. psl.*) complex. The *C. psl.* complex belongs to Zygnematophyceae, which is closely related to land plants (Wickett et al. 2014; fig. 1a). In *Closterium*, the haploid cells proliferate through vegetative reproduction, while under stressful conditions, two cells form a dormant zygospore to reproduce sexually (fig. 1b and c). In the *C. psl.* complex, two modes of zygospore formation have been reported: homothallism and heterothallism (Tsuchikane and Sekimoto 2019; Ohtaka and Sekimoto 2023). The homothallic strains can form zygospores within clonal cells (fig. 1b), whereas the heterothallic strains can form zygospores between cells of two different mating types: plus (mt^+^) and minus (mt^−^; fig. 1c). In heterothallic strains, mt^+^ and mt^−^ recognize each other through the sex pheromones and then form zygotes (Sekimoto et al. 1994; Nojiri et al. 1995). Heterothallic strains have been further classified into reproductively isolated mating groups. Members of the different mating groups cannot mate with each other, except in the cases where reproductive isolation between groups II-A and II-B and between groups G and I-E is incomplete and asymmetric (Watanabe and Ichimura 1978; Tsuchikane et al. 2008; Tsuchikane et al. 2018a). Quite recently, genomes of both mt^+^ (NIES-67) and mt^−^ (NIES-68) and the major genetic factor responsible for mating-type determination have been reported in group I-E (Sekimoto et al. 2023).

**Fig. 1.**
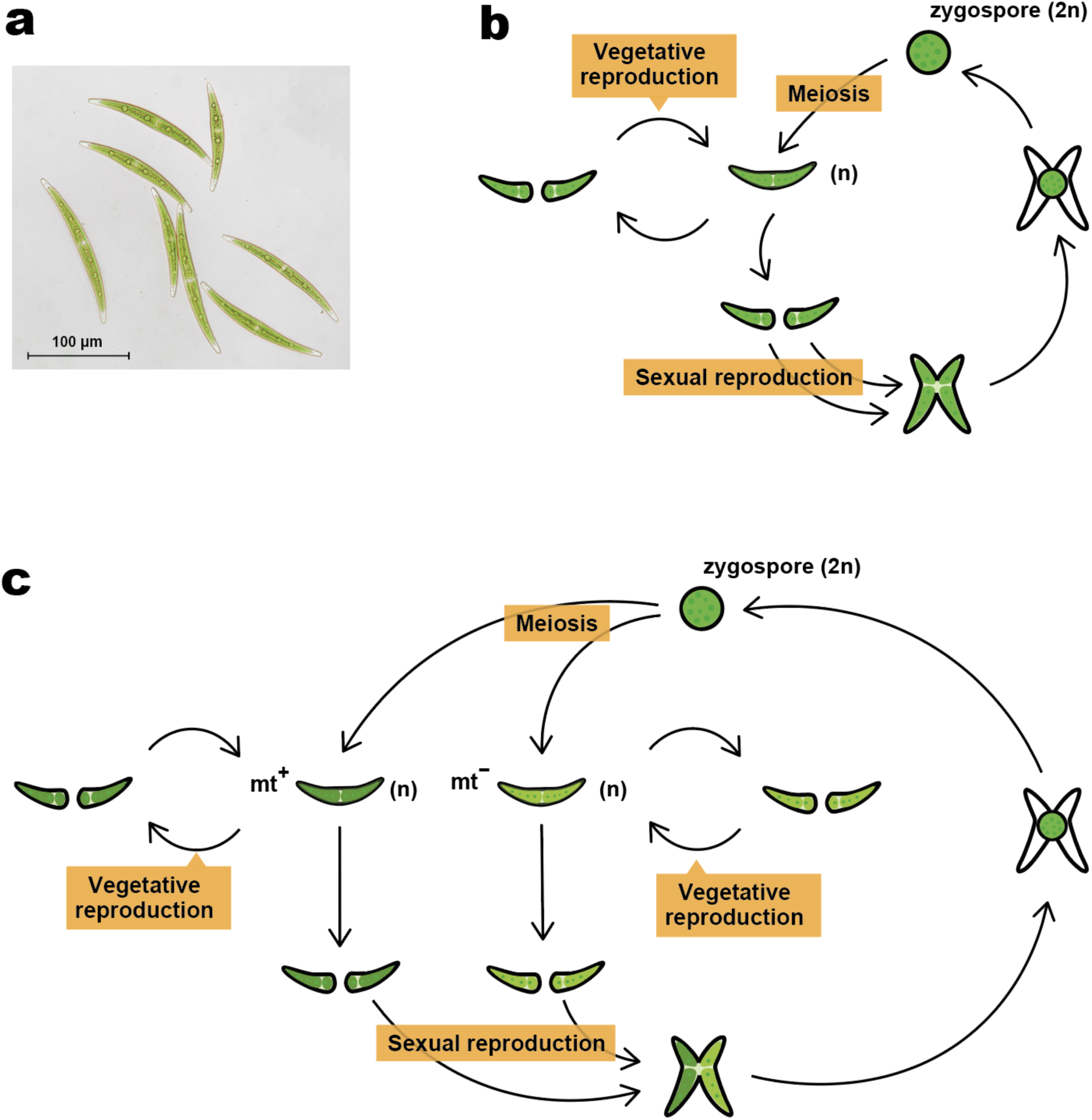
The life cycle of the *Closterium peracerosum-strigosum-littorale* (*C. psl.*) complex. (a) A photograph of vegetative cells of the NIES-65 strain in the *C. psl.* complex. (b,c) Schematic illustrations of the life cycle of homothallic (b) and heterothallic (c) strains in the *C. psl.* complex.

In this study, we identified twofold genome size variation among these natural strains using a short-read-based *k*-mer analysis. Flow cytometry of F_1_ hybrids, cells germinated from zygospores obtained by crossing, revealed that genome size variation is heritable and not significantly associated with mating types. *De novo* assembled genomes of six strains confirmed the genome size difference and revealed that CNV among strains was most strongly associated with genome size variation. Transcriptome analysis revealed that dosage compensation occurs in about 30% of genes with CNV. Thus, the expression level does not necessarily increase proportionally with the gene copy number in those genes. We found that specific gene categories, such as those involved in translation, are enriched in the dosage-compensated genes. The *C. psl.* complex serves as a new model system, paving the way for studying genome size evolution and how organisms functionally mitigate the copy number changes.

## Results

### Phylogenetic Relationship and Genome Size Variation in the *Closterium psl.* Complex

We first obtained short-read sequence data from 22 strains of the *C. psl.* complex to perform a phylogenetic analysis and estimate the genome size (supplementary table S1). As a reference for the short-read mapping, we also generated a *de novo* assembled sequence of the NIES-4552 strain with a total length of 438 Mb and N50 of 952 kb (supplementary table S2). We obtained 79,377 SNPs by mapping the short reads of 22 strains and constructed a phylogenetic tree using the maximum likelihood method. The tree showed that each of the five mating groups of heterothallic strains (II-A, II-B, II-C, I-E, and G) formed a distinct clade. In comparison, homothallic strains were separated into three groups, indicating multiple transitions of mating systems between heterothallism and homothallism (fig. 2).

**Fig. 2.**
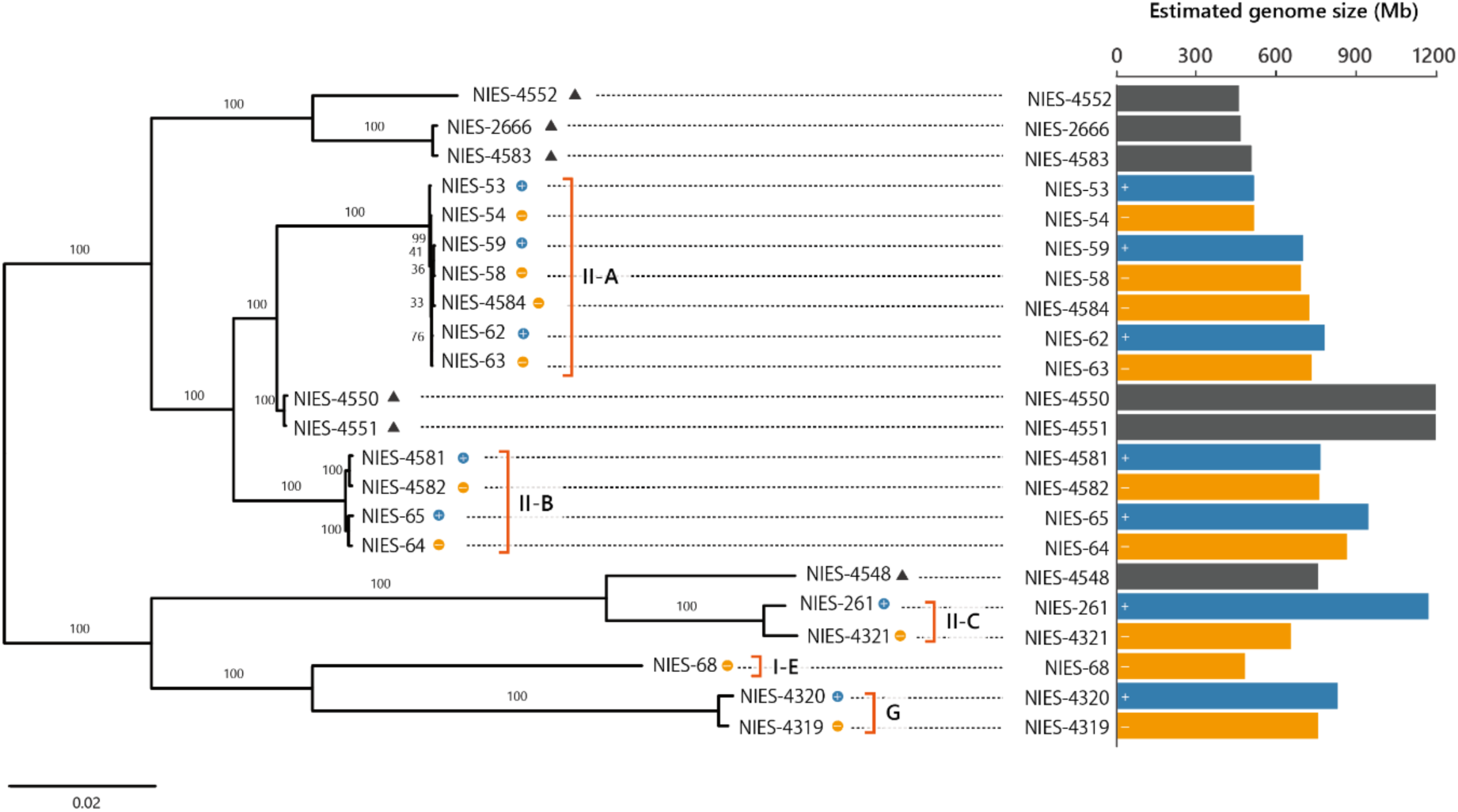
Phylogenetic relationship and estimated genome size variation in the 22 strains of the *Closterium peracerosum-strigosum-littorale* complex. The tree was constructed using the maximum likelihood method based on 79,377 SNPs obtained through mapping to the NIES-4552 strain as a reference. *k*-mer analyses estimated the genome size. II-A, II-B, II-C, I-E, and G indicate mating groups. Black triangles, blue points, and orange points indicate homothallic, heterothallic (mt^+^), and heterothallic (mt^−^) strains, respectively. The bars show the genome size of the respective strains. Mating types (mt^+^ and mt^−^) are defined for each mating group and may not be comparable across II-A, II-B, II-C, I-E, and G, as different mating groups cannot reproduce with each other.

We further exploited short-read sequences to estimate the genome size of each strain using *k*-mer analysis. To estimate genome size, including high copy sequences, we calculated the total number of all *k*-mers without an upper limit and subtracted the peaks of organelle genomes from it (see Materials and Methods for details). The estimated genome size showed a more than twofold variation among strains, ranging from 454 to 1,188 Mb (fig. 2, supplementary table S3). Notably, genome size showed 16.2% and 24.3% differences even within the heterothallic mating groups II-A and II-B, respectively.

Mating types (mt^+^ and mt^−^) and genome size (*p* = 0.775 for the group II-A; *p* = 0.723 for the group II-B; Student’s *t*-test) were not associated significantly, suggesting that genome size variation is not solely explained by structural variation in sex-linked genomic regions. This finding is consistent with a report on a limited sex-linked region in group I-E (Sekimoto et al. 2023). After excluding high copy sequences, the association between genome size variation and mating types was still nonsignificant (*p* = 0.827 for the group II-A; *p* = 0.569 for the group II-B; Student’s *t*-test; supplementary table S3, supplementary fig. S1, see Materials and Methods for details).

### Inheritance of Genome Size Variation

Since we found extensive genome size variation even within the same mating groups, we next investigated whether and how genome size is inherited to the next generation. We established F_1_ hybrids between two heterothallic strains, NIES-64 and NIES-65, and estimated genome size of 18 F_1_ hybrids by flow cytometry of isolated nuclei stained with propidium iodide. Each strain was measured at least three times, and we found that genome size is heritable (51.9% of total genome size variance was due to the variance between F_1_ lines; *p* = 1.36 × 10^−6^; ANOVA) and that the median of genome size of all F_1_ lines was in between those of the two parental strains (fig. 3a). We also found that genome size variation was not significantly deviated from the unimodal distribution (*p* = 0.566; unimodality test; Ameijeiras-Alonso et al. 2021) and not significantly associated with mating types (*p* = 0.173; Student’s *t*-test; fig. 3b). When F_1_ lines were narrowed down to those with the minimum number of subculturing (three times, corresponding to about 50–200 generations of vegetative reproduction) to minimize the effect of vegetative reproduction within each line (supplementary fig. S2), we still observed about 130 Mb difference (∼90% of parental difference) between F_1_ lines, indicating that genome size variation is inherited during the sexual process at least partially. Together with the results of *k*-mer analysis, these findings overall suggest that the genome size variation between NIES-64 and NIES-65 is not due to the structural variation in specific genomic fragments such as sex-linked regions but rather due to structural variation in multiple genomic regions.

**Fig. 3.**
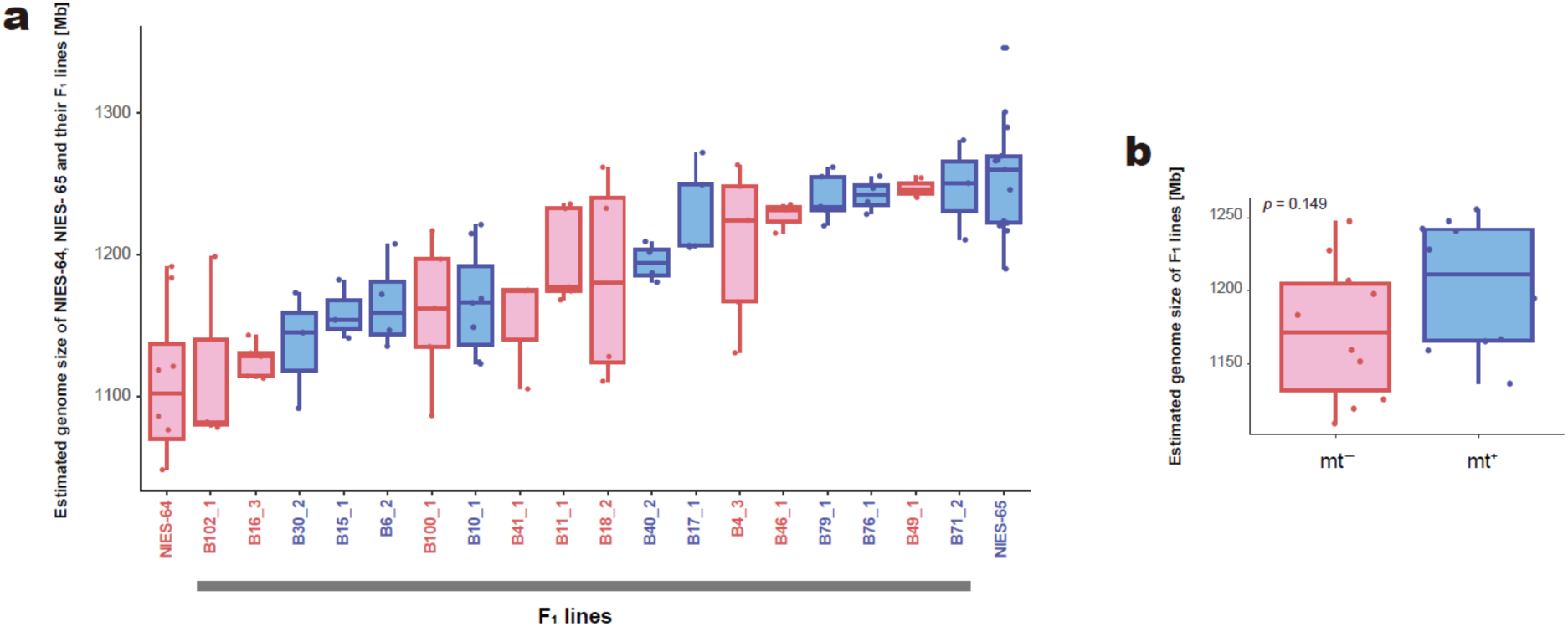
Genome size variation of F_1_ lines estimated by flow cytometry. (a) Genome size of each F_1_ line. Red and blue bars indicate mating types of F_1_ lines, mt^−^ and mt^+^, respectively. (b) Genome size was not significantly different between mating types (*p* = 0.149; *t*-test). Each dot shows the average genome size of each F_1_ line. Bars and boxes represent the median and the interquartile range, respectively. Whiskers extend to 1.5 times the interquartile range.

### Genome Size Variation is Explained Largely by Gene Duplication

To determine the factors contributing to genome size variation, we focused on six strains (NIES-4552, NIES-4550, NIES-64, NIES-65, NIES-53, and NIES-54) that differ greatly in genome size. We performed long-read whole-genome sequencing and *de novo* assembly for these six strains (supplementary table S2). As a result, the total length of the *de novo* assembled sequence varied from 438 (NIES-4552) to 975 Mb (NIES-4550), which was consistent with the estimates from the *k*-mer analyses (correlation coefficient = 0.960; *p* = 0.00238; supplementary fig. S3a). We also obtained genome size estimates based on flow cytometry (Tsuchikane et al. 2023), which were significantly correlated with both the total length of the *de novo* assembled sequences and the estimates from the *k*-mer analyses (correlation coefficients: 0.974 [*p* = 0.00103] and 0.879 [*p* = 7.28 × 10^−6^], respectively; supplementary fig. S3b and c). We also compared the nuclear size between two strain NIES-4550 (estimated genome size: 975 Mb) and NIES-4552 (estimated genome size: 438 Mb), finding that the nuclear size of the NIES-4550 was significantly larger than that of the NIES-4552 (*p* = 7.34 × 10^−6^; Student’s *t*-test), consistent with the genome size estimates (supplementary fig. S4).

We then performed gene annotation for each genome to compare gene contents among the strains. Overall, the proportion of detected proteins of Benchmarking Universal Single-Copy Orthologs (BUSCO) genes of Viridiplantae was ˃85% in all strains (supplementary table S2). In comparison, “complete” was ˂70% in NIES-53 and NIES-54 and ˃83% in the other four strains, showing high levels of completion of assembly and gene annotation in the four strains and moderate draft levels in NIES-53 and NIES-54 (supplementary table S2). We identified ortholog groups (OGs) with the SonicParanoid software (Cosentino and Iwasaki 2019). We then determined the copy number of each OG in the six strains. Focusing on the 5,400 OGs shared by all strains, we found that strains with larger genome sizes tended to have more duplicated genes (*p* = 0.0315; Pearson’s product-moment correlation coefficient; fig. 4a and b). Notably, more than half of the OGs were duplicated in the strains NIES-65 and NIES-4550. Given that analyzed samples were obtained during the haploid phase, this result suggests the presence of aneuploidy or segmental duplications as indicated by the bimodal frequency distributions in the *k*-mer analysis (supplementary fig. S5). These results further suggest that CNV largely contributes to genome size variation among strains.

**Fig. 4.**
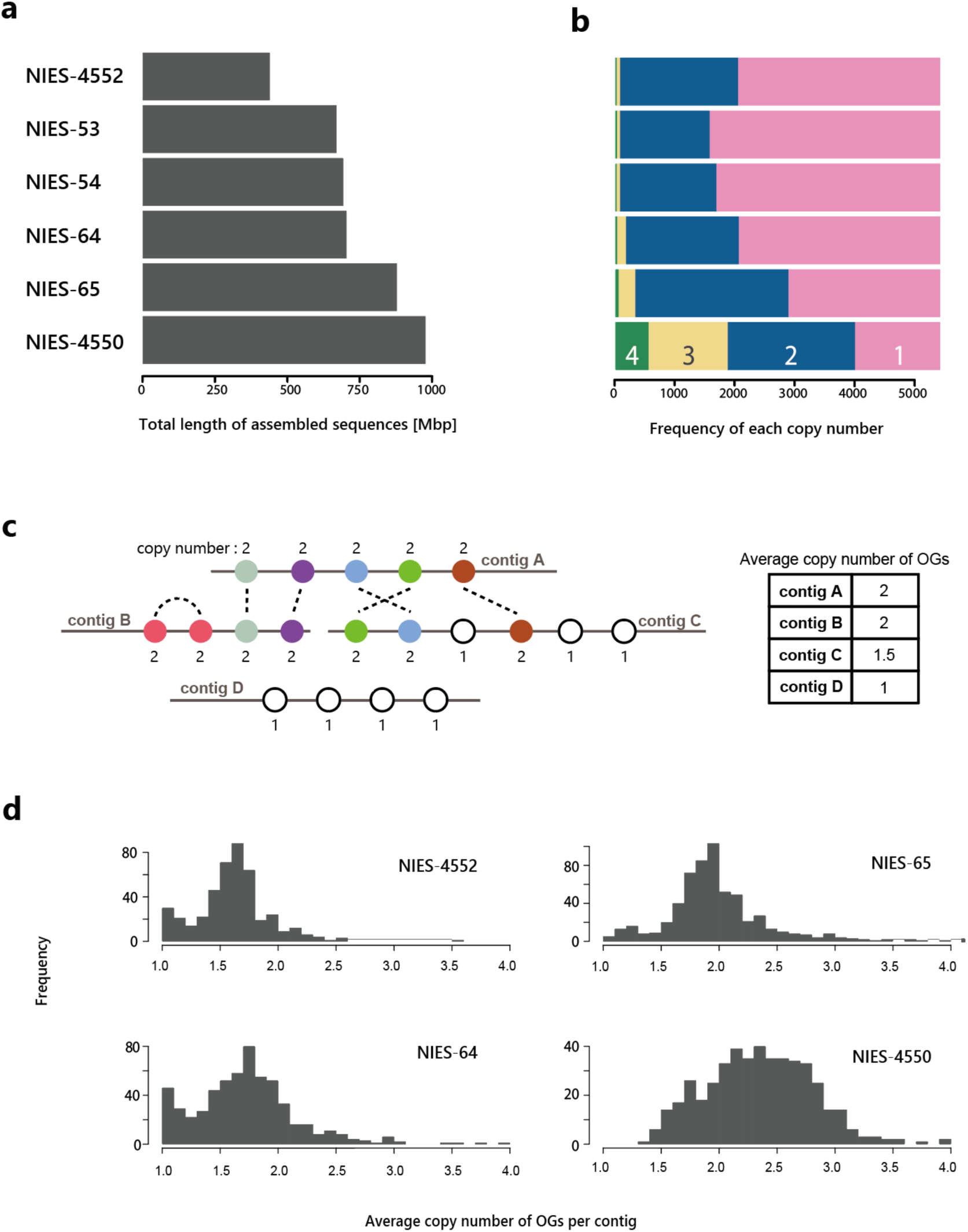
Total genome size and copy number variation of *de novo* assembled genomes. (a) The total length of *de novo* assembled sequences. (b) Copy numbers of ortholog groups (OGs) shared by six *de novo* assembled genomes. The numbers in the bar indicate the copy number. We defined the copy number as the number of genes assigned to the identical OG. (c) A schematic diagram illustrating how to calculate the average copy number of OGs per contig. (d) Distribution of the average copy number of the OGs per contig of four strains with high-quality genomes.

We next investigated if other genome characteristics contribute to genome size variation: specifically, the number of small copy number genes (˂5 copies), the number of high copy number genes (˃9 copies), the average length of genes, introns and intergenic regions, the average number of introns, and the proportion of intergenic regions (table 1). Because of the relatively low contiguity of the NIES-53 and NIES-54 draft genomes (N50 = 51 and 40 kb, respectively), we used the other four high-quality genomes (NIES-4552, NIES-4550, NIES-64, and NIES-65) for these analyses. Among the characteristics analyzed, the number of small copy number genes, multiple-copy genes, and tandem repeat genes was significantly correlated with genome size variation (table 1). The distribution of the average copy number of the OGs per contig showed peaks from 1.5 to 3 in all strains, suggesting the presence of duplicated genes in most contigs (fig. 4c and d). The strains NIES-4552 and NIES-64 also had peaks at an average copy number of 1, indicating the presence of contigs without duplicated genes. The dot plots of self-alignments also supported the observation that there are of duplicated genes distributed across the genome (supplementary fig. S6). The number of tandem repeat sequences, defined by duplicated genes on the same contig, correlated with genome size; however, their proportions were less than 1% in any strain and their contributions to the genome size variation were considered minor. The proportion of repeat sequences varied among strains (47.7% for NIES-4552, 64.4% for NIES-64), but the proportion did not correlate with genome size, indicating that repeated sequences were not the primary factor contributing to genome size variation (table 1, supplementary table S5). We note that the proportion of repeat sequences also did not correlate with flow cytometry-based genome size estimates (*p* = 0.575). Overall, these results suggest that numerous gene duplications are the main factor contributing to genome size variation. Since the distribution of duplicated genes was found to be genome-wide, the variation would have been generated through whole-genome duplications followed by subsequent deletions or numbers of segmental duplications throughout the genomes.

**Table 1.** Comparison of key features of four high-quality genomes.

|  | NIES-4552 | NIES-64 | NIES-65 | NIES-4550 | *p* values | Correlation coefficient |
| --- | --- | --- | --- | --- | --- | --- |
| Total length | 438 Mb | 703 Mb | 876 Mb | 975 Mb | - | - |
| Annotated genes | 27,822 | 34,905 | 40,587 | 52,529 | 0.061 | 0.939 |
| Small copy number genes (<5) | 17,026 | 22,000 | 24,321 | 29,234 | 0.028 | 0.972 |
| High copy number genes (>9) | 1,253 | 2,637 | 3,770 | 2,414 | 0.281 | 0.719 |
| Single-copy genes<br>(no. ortholog group) | 9,615 | 12,280 | 10,654 | 9,789 | 0.939 | 0.061 |
| Multiple-copy genes<br>(no. ortholog group) | 3,662 | 13,363 | 18,477 | 23,631 | 0.003 | 0.997 |
| Tandem repeat genes<br>(no. ortholog group) | 105 | 124 | 141 | 159 | 0.019 | 0.981 |
| Proportion of tandem repeat genes<br>(no. ortholog group) | 0.791% | 0.484% | 0.484% | 0.476% | 0.113 | -0.887 |
| Mean length of transcripts | 2,014 bp | 2,000 bp | 2,173 bp | 2,114 bp | 0.251 | 0.749 |
| Proportion of repeated elements | 47.2% | 64.4% | 62.6% | 53.7% | 0.528 | 0.472 |
| Mean length of introns | 548.9 bp | 488.2 bp | 485.4 bp | 554.1 bp | 0.840 | -0.160 |
| No. introns per gene | 5.7 | 5.2 | 5.0 | 5.6 | 0.615 | -0.385 |
| Mean length<br>of intergenic region | 10,237 bp | 14,306 bp | 15,935 bp | 13,070 bp | 0.300 | 0.701 |
| Proportion<br>of intergenic region | 67.3% | 77.4% | 78.7% | 72.0% | 0.440 | 0.560 |

The *p* values and correlation coefficient indicate the result of Pearson's product-moment correlation between the genome size and each feature.

### Independent Gene Duplications and Hybridizations

We next addressed the origin of duplicated genes. We first focused on the homothallic NIES-4550 strain due to its largest genome size and the highest gene copy number among the studied strains. Short-read genomic sequences of heterothallic strains of the mating groups II-A (NIES-58 and NIES-59) and II-B (NIES-64 and NIES-65) were mapped to the assembled genome of NIES-4550. We found that the coverage depth of reads from groups II-A and II-B showed a complementary pattern: genes with high coverage of group II-A reads tended to have low coverage of group II-B reads and vice versa. This result highlights the heterogeneous nature of the genome of the NIES-4550 strain (fig. 5a).

**Fig. 5.**
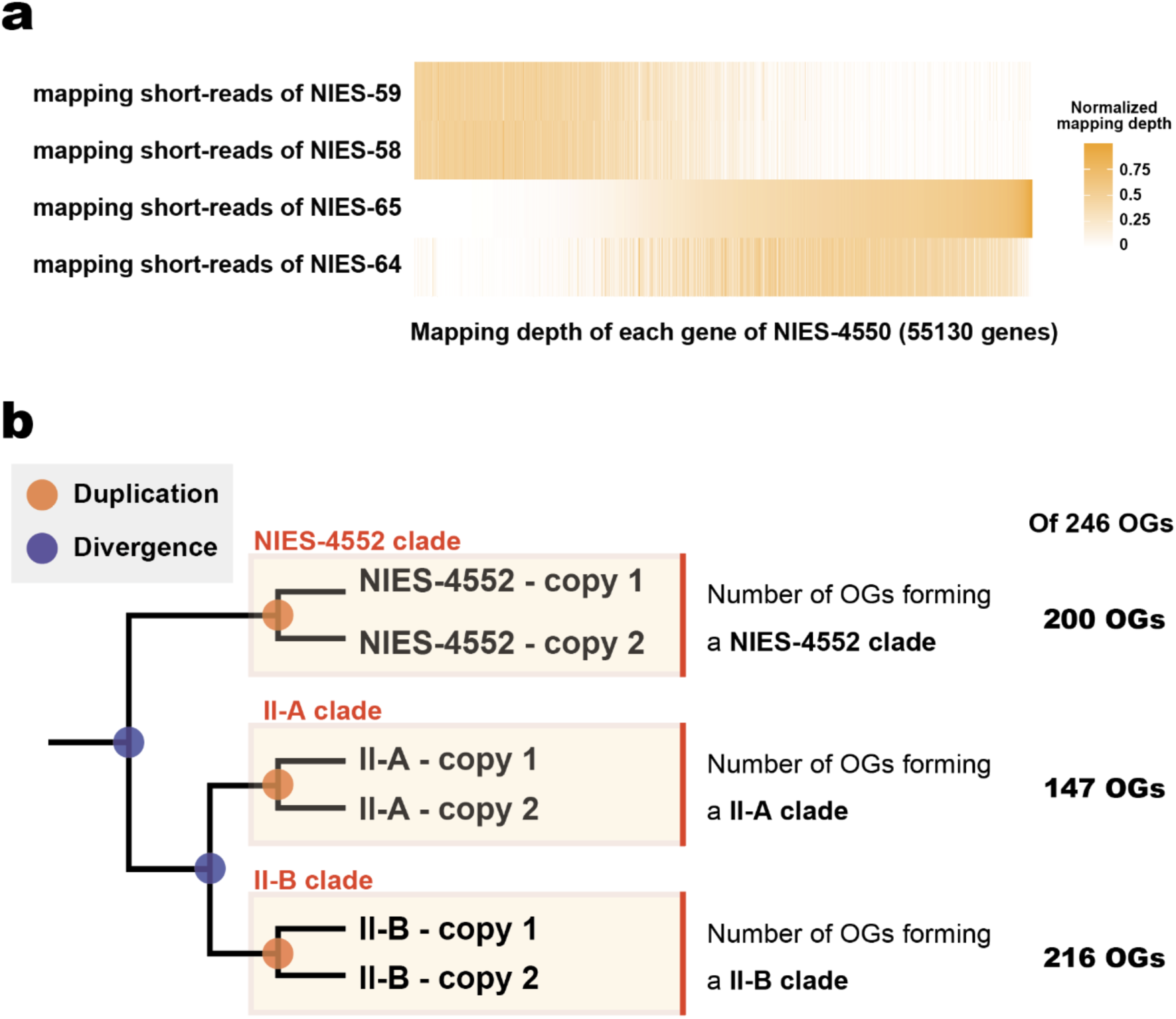
The origins of duplicated genes found in the group II-A, the group II-B, the NIES-4550 strain, and the NIES-4552 strain. (a) The mapping depth of short reads from groups II-A (NIES-58 and NIES-59) and II-B (NIES-64 and NIES-65) to the assembled genome of NIES-4550. (b) Clarifying the origins of duplicated genes found in the group II-A, the group II-B, and the NIES-4552 strain. If duplications occurred independently in each strain, gene copies from the same strains should form a clade. The origins of duplicated genes of the 246 ortholog groups (OGs) that have two to four copies in the group II-A, the group II-B, and the NIES-4552 strain. Each group formed a clade with high bootstrap support (≥ 70%) in 147–216 OGs. Note that this is a schematic phylogenetic tree, and three clades are shown in a single figure for simplicity. In the actual analysis, the monophyly of three groups was investigated separately.

The ortholog-based phylogenetic approach also supported the heterogeneous nature of the NIES-4550 strain. Of the 989 OGs that were duplicated only in the NIES-4550 strain, 351 OGs showed the pattern that a gene of NIES-4550 clustered with the mating group II-A and the other gene clustered with the mating group II-B (bootstrap ≥ 70; supplementary table S6). This result suggests that the genome of the NIES-4550 strain contains genes closely related to the mating groups II-A and II-B. On the other hand, the NIES-4550 genome contained 4,250 genes that are distantly related from both the mating groups II-A and II-B by evolutionary distance analyses, suggesting that at least another ancestor contributed to the NIES-4550 genome in addition to the mating groups II-A and II-B (see Supplementary Note for details). We found at least 103 contigs that include genes with different presumptive origins, suggesting that recombination between sequences of different origins occurred (supplementary fig. S7a). The physical connection between genomic regions of different presumptive origins was confirmed experimentally by PCR amplification and Sanger sequencing of recombination borders (supplementary figs. S7b and c and S8, see Supplementary Note for details). In summary, the heterogeneity of the NIES-4550 genome reflects its hybrid origin with subsequent rearrangements, including recombination between genes of different origins.

We constructed a phylogenetic tree for each OG to determine the origin of the duplicated genes found in the other five strains. Assuming that the duplicated genes originated from a gene duplication in a common ancestor, genes in a strain should form separate clades containing genes from both strain. If the duplication occurred in each strain after their separation, genes from a strain should form a clade without genes from the other strain, although concerted evolution by gene conversion may also result in a similar pattern (fig. 5b). We did not assume ancestral polymorphisms or crossbreeding between strains; thus, we did not distinguish between strains belonging to the same mating groups.

Focusing on the 246 OGs that have two to four copies in all five strains other than the NIES-4550 strain, we examined whether each strain or the mating group forms a clade. By classifying the tree topologies, we found that the group II-A, the group II-B, and the NIES-4552 strain formed a clade with high bootstrap support (≥ 70%) in 147, 216, and 200 OG trees, respectively (fig. 5b, supplementary table S7). Most of the OG trees (239 OGs) was found to form clades either in the group II-A, the group II-B, or the NIES-4552 strain with high bootstrap support (≥ 70%; fig. 5b, supplementary table S7). These results indicate that at least a part of duplicated genes found in the group II-A, the group II-B, and the NIES-4552 strain originated from recent duplications within each group.

### Correlation Between Expression Level and Gene Dosage

Given that the genomes of the *C. psl.* complex have high variability in gene copy numbers among strains, a compensating mechanism might be buffering gene expression, thereby tolerating the copy number changes. To investigate the relationship between copy number and gene expression, we performed RNA-seq analyses in the strains NIES-64 and NIES-65 at five different stages: during vegetative reproduction at morning (L), during vegetative reproduction at night (D), under a nitrogen-depleted condition (M), and in conditioned mating-inducing medium for 8 h (C8h) and 24 h (C24h; fig. 1; supplementary table S8, see Materials and Methods for details).

Between the two strains, 31.6% of OGs showed copy number difference based on an orthologous grouping using the SonicParanoid software (Cosentino and Iwasaki 2019; fig. 6a). The high fraction of genes showing copy number difference was also confirmed by the DNA depth data. The DNA depth log-ratio of each OG was defined as the difference in log_2_-transformed read counts normalized for the genome (NIES-65 – NIES-64). We changed the criteria of the DNA depth log-ratio for the definition of CNV depending on the total DNA depth, i.e., the sum of normalized log_2_-transformed read counts of NIES-64 and NIES-65. This is because the values of non-zero peaks, indicating CNV, differed depending on the total DNA depth (fig. 6b, supplementary fig. S9, see Materials and Methods for detail). Based on the criteria, we found that 36.2% of OGs showed the high DNA depth log-ratio between NIES-64 and NIES-65 (fig. 6b).

**Fig. 6.**
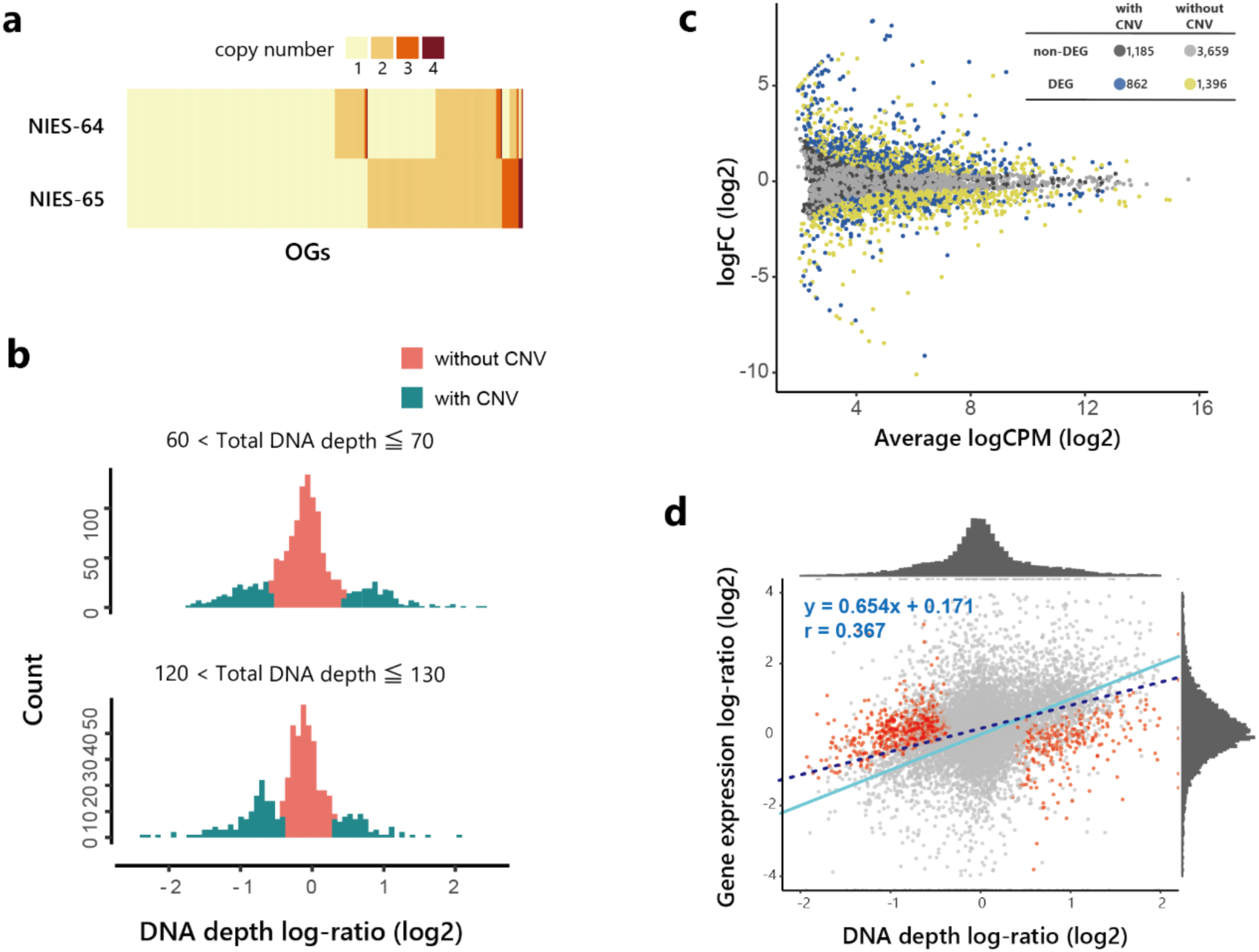
Correlations between expression levels and gene dosages. (a) Copy number difference of ortholog groups (OGs) between the NIES-64 and NIES-65 strains. We defined the copy number as the number of genes assigned to the identical OG. OGs with more than four copies are not shown. (b) Frequency of DNA depth log-ratio for each OG (see supplementary fig. S9 for other ranges of a total DNA depth). (c) An MA plot of gene expression difference between the NIES-64 and NIES-65 strains. Expression data from the stage L were used (see supplementary fig. S10 for other stages). The number of genes and color in each category, differentially expressed genes (DEGs) or not, and with or without copy number variation (CNV), is shown in the upper right table. The genes with CNV were defined based on the DNA depth log-ratio (see Materials and Methods for details). (d) The expression level and gene dosage difference between the NIES-64 and NIES-65 strains. Expression data from the stage L were used (see supplementary fig. S11 for other stages). A light blue line shows the expected relationship representing proportional increases in expression difference to dosage difference, while a blue dashed line represents the simple regression. Dosage-compensated genes are shown in red.

We investigated the relationship between the expression level and the copy number between two strains, NIES-64 and NIES-65. We found that 31.8–39.5% OGs showed significantly different expressions between the two strains (false discovery rate (FDR) ˂ 0.05). For the following analysis, the definition of copy number was based on the DNA depth log-ratio. In every stage, the presence of CNV was associated with the difference in expression levels: genes with CNV were significantly enriched in the set of differentially expressed genes (DEGs; fold enrichment ˃ 1.24, *p* ˂ 2.2 × 10^−16^; Fisher’s exact test; fig. 6c, supplementary fig. S10). We also found that expression level differences and gene dosage differences were positively correlated (*p* ˂ 0.01, correlation coefficient ˃ 0.344; fig. 6d, supplementary fig. S11), but the slopes (regression coefficient) were 0.654–0.683, significantly smaller than 1 (*p* ˂ 0.001); the expression level overall responded to gene dosage, but less than proportionally. In addition, the gene-specific analysis revealed that 30.6–39.7% of the OGs with CNV showed significantly smaller expression level differences than expected from an assumption that expression level is proportional to the gene dosage (FDR ˂ 0.05; fig. 6d, supplementary fig. S11). We found that 6.87–13.3% genes showed significantly higher expression level difference than proportional to gene dosage (FDR ˂ 0.05), which was defined as genes with a positive dosage response. Although the frequency of genes with a positive dosage response was generally low, the frequency was relatively high in the sexual reproductive stage, suggesting that these genes reflect mating type-specific differences in expression (7.28% in L, 6.87% in D, 9.78% in M, 11.0% in C8h, and 13.3% in C24h). Furthermore, among the genes for which the expression level difference was smaller than proportional to the gene dosage, 18.4–29.9% of genes had a significantly higher expression in the low copy number strain than in the high copy number strain (FDR ˂ 0.05; “inverse effect” by Guo and Birchler 1994). We defined dosage-compensated genes as genes not showing the inverse effect (FDR ˃ 0.1) but showing significantly smaller expression level differences than proportional to gene dosage (following Zhang et al. 2017), identifying 24.9–29.4% of genes with CNV as dosage-compensated genes.

### Gene Function and Expression Level Variation

Since we found different gene expression patterns associated with gene dosage, we investigated which biological functions might be related to dosage sensitivity of gene expression and CNV between NIES-64 and NIES-65. We performed gene ontology (GO) enrichment analyses for dosage-compensated genes. We found in total 27 significantly enriched GO terms through the analyses of two or more stages (fig. 7). In the set of dosage-compensated genes, we found the enrichment of “translation” (GO:0006412), “structural constituent of ribosome” (GO:0003735), “mRNA binding” (GO:0003729), “mRNA splicing, via spliceosome” (GO:0000398), “Golgi membrane” (GO:0000139), and “trans-Golgi network” (GO:0005802; fig. 7, supplementary table S9–S13). The results of enrichments suggest that the expression levels of genes involved in translation and the trans-Golgi network, tended to be robust to gene dosage changes. Interestingly, we also found the enrichment of “mRNA splicing, via spliceosome” (GO:0000398), “mRNA binding” (GO:0003729), “regulation of mRNA stability” (GO:0043488), “ribosome biogenesis” (GO:0042254) and “structural constituent of ribosome” (GO:0003735) in the gene set that does not show CNV between NIES-64 and NIES-65 (supplementary table S14). These results overall suggest that genes involved in translation are less likely to have CNV, and that even when CNV is observed, expression level difference is less than proportional to copy number difference.

**Fig. 7.**
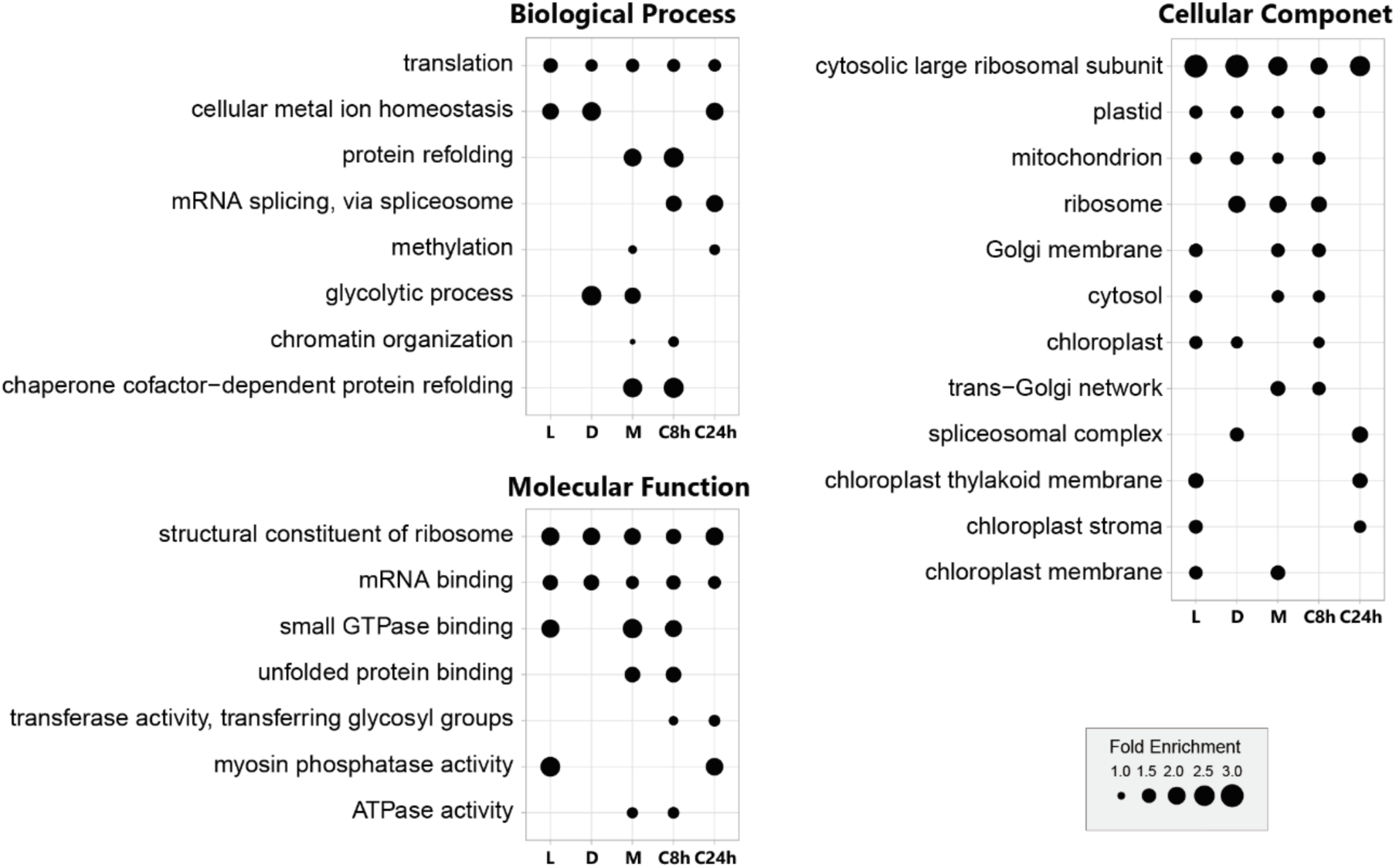
Gene ontology (GO) enrichment analysis of dosage-compensated genes. Among the GO terms including more than nine genes, those with enrichment of dosage-compensated genes in at least two stages are shown (*p* ˂ 0.05; Fisher’s exact test with the ‘weight01’ algorithm in the R package topGO, Alexa and Rahnenführer 2009). The dot size indicates the fold enrichment. Each column shows results during each stage: during vegetative reproduction in light (L), during vegetative reproduction in dark (D), under a nitrogen-depleted condition (M), and in the mating-induced conditioned medium for 8 h (C8h) and 24 h (C24h).

We also found that the genes without CNV were overrepresented in the gene set that showed constitutive expression along five different stages (*p* = 3.63 × 10^−14^, supplementary table S15). Conversely, genes with CNV were overrepresented in genes which were not expressed in one to four stages (*p* = 0.0201, supplementary table S15), indicating that genes with stage-specific expression are more prone to change gene dosage by duplication or deletion.

## Discussion

Although genome size variation within or among closely related species is common, few studies so far have addressed the genomic factors contributing to the variation and its consequences for gene function (Šmarda et al. 2010; Long et al. 2013; Huang et al. 2014; Sun et al. 2018). Here we revealed the more than twofold heritable genome size variation among closely related strains in the *C. psl.* complex was largely attributable to genome-wide CNV rather than the number of tandem repeat sequences or the proportion of repeat sequences (table 1, figs. 2 and 4). Notably, more than 30% of OGs showed CNV even between genetically close, sympatric strains that can mate with each other, namely NIES-64 and NIES-65 (fig. 6a and b). Here we note a few caveats for these findings. First, the reliability of these analyses depends on the quality of *de novo* assembly, because the genome size, the number of repeat sequences, and the fraction of CNV are all strongly influenced by the quality of *de novo* assembly. Nonetheless, flow cytometry-based genome size estimates showed significant correlations with both the total length of the *de novo* assembled sequences and the estimates from the *k*-mer analyses. Since the flow cytometry-based estimation is independent from sequence artifacts, this result supports the reliability of our sequence-based genome size estimates, although the contribution of repeat sequences missed in assembly is not fully ruled out (supplementary fig. S3). Second, while we discuss the inheritance of genome size variation, variation found among natural strains should include both mutations occurred during sexual reproduction and those accumulated during vegetative reproduction, which cannot be distinguished by our approach of comparing genome sequences. Yet, differences among F_1_ lines were still observed when narrowing down the samples to reduce the effect of vegetative reproduction (supplementary fig. S2). Furthermore, the flow cytometry analysis of F_1_ hybrids revealed that 51.9% of total genome size variance was due to the variance among F_1_ lines and that the median of genome size of all F_1_ lines was in between those of the two parental strains. In summary, although we acknowledge these caveats and limitations, independent lines of evidence from *de novo* assembly, *k*-mer-based analysis, and flow cytometry analysis consistently suggest that genome size variation is largely attributable to genome-wide CNV and is inherited during the sexual process at least partially.

CNV has been reported in several species: copy number variable regions cover 12% of the human genome (Redon et al. 2006), and 8% of the regions of the *Drosophila melanogaster* genome were estimated to contain CNVs (Dopman and Hartl 2007). The fraction of OGs showing CNV found in the *C. psl.* complex (about 30%) is strikingly high compared with these reports. We note that this would still be underestimated as the CNV within species because here we compared a single pair of genetically close, sympatric strains. Recently, the cat flea, exhibiting a broad range of genome size variation, was also shown to have a genome with inordinate CNV (∼38% of proteins; Driscoll et al. 2020). The fraction of CNV and genome size variation appears to be highly diverse among lineages, and accumulating knowledge in a wide range of organisms will allow us to understand which factors determine those variations across lineages.

Why is such a high level of CNV harbored in the *C. psl*. complex, despite the possible detrimental effects of CNVs (Torres et al. 2007; International Schizophrenia Consortium 2008; Cooper et al. 2011; Dephoure et al. 2014)? Our study suggests that possible detrimental effects could be diminished in multiple ways. CNV is prevalent in the *C. psl*. complex, and genes showing CNV are nonrandom in terms of gene functions. Specifically, we found that (1) genes involved in basic biological functions were less likely to have CNV, (2) CNV was more often observed in the gene set with stage-specific expression, and (3) genes without CNV were overrepresented in the genes with constitutive expression. A large-scale analysis across angiosperms revealed that likelihood of gene duplications differs depending on gene functions, such that single-copy genes are enriched for “essential” genes that are required for early development and survival. By contrast, “conditional” genes that respond to biotic and abiotic stress are more prone to duplicate (Li et al. 2016). Similar gene functions and duplicability patterns are also demonstrated in fungi (Wapinski et al. 2007).

Our transcriptome analyses also revealed that the expression level does not increase proportionally with the gene copy number, suggesting the presence of a compensating mechanism that buffers gene expression (fig. 6d, supplementary fig. S11). Such a mechanism may tolerate the gene dosage changes, possibly allowing CNVs in the genomes. The GO analysis demonstrated that the responsiveness to gene dosage differs depending on gene function: genes involved in basic biological functions, such as translation, especially ribosomal proteins, were enriched in the dosage-compensated genes (fig. 7). Enrichment of those gene sets involved in translation was also observed in the genes subject to dosage compensation in aneuploidy strains of *Saccharomyces cerevisiae* (Hose et al. 2015), and dosage compensation of ribosome-related genes has also been reported in other organisms, including flies, wheat, and humans (Hose et al. 2015; Lee et al. 2016; Zhang et al. 2017; Hwang et al. 2021; Schukken and Sheltzer 2022). We note that results of gene set enrichment analyses should be interpreted with caution because of its potential bias due to the heterogeneity of the source data (Gaudet and Dessimoz 2016). There are a few experimental supports for dosage compensation of translation-related genes. Artificial changes in the copy number of rRNA and ribosomal proteins did not alter their expression proportional to the copy number (Pearson et al. 1982; Lopez et al. 2021), possibly by the coordinate regulation of ribosomal proteins and rRNA to keep dosage balance (Warner 1999). An analysis of the hybrid genomes of *Trichosporon* revealed that the gene-loss rate among protein complexes involved in translation was significantly higher than that among other complexes (Sriswasdi et al. 2016). Our study suggests that dosage compensation, particularly in translation, is a common phenomenon in many organisms. In summary, gene duplication and expression changes in response to gene duplication are nonrandom in terms of gene functions, possibly contributing to the maintenance of the high level of CNV in association with extensive genome size variation in the *C. psl*. complex. We also note that a recent study reported a difference of chromosome number in association with genome size difference even within the same mating group, suggesting a mechanism tolerating chromosomal rearrangements during meiosis in the *C. psl*. complex (Tsuchikane et al. 2023). This organism could serve as a new model system, paving the way for studying the genome size evolution and how organisms functionally mitigate the potentially pernicious effects of copy number changes.

## Materials and Methods

### Studied Strains

We used in total 22 strains of the *C. psl.* complex (supplementary table S1), namely 6 homothallic strains and 16 heterothallic strains (including 6 mt^+^ strains were and 10 mt^−^ strains). These strains were obtained from the National Institute for Environmental Studies (NIES), Environment Agency (Ibaraki, Japan). Mating group I-E is denoted by IB in the NIES.

### Culture and Mating Conditions

All strains were cultured in a nitrogen-supplemented medium (CA medium, Kasai et al. 2004) at 21°C in a 16 h light/8 h dark cycle. To induce sexual reproduction, cells vegetatively grown for ten days were washed three times with a nitrogen-depleted mating-inducing medium (MI medium, Ichimura 1971). Then, one or two strains were mixed and incubated in an MI medium (10,000 cells/mL) under 24 h continuous light.

### DNA and RNA Extraction

DNA was isolated from cells vegetatively grown for two weeks. For NIES-53 and NIES-54, we extracted DNA using NucleoBond HMW DNA (Takara Biotechnology, Japan) and eliminated the short fragments using the Short Read Eliminator Kit (PacBio). Following Shure et al. (1983), we extracted DNA for other strains and purified it using a Blood and Cell Culture DNA mini kit (Qiagen). Finally, total RNA was extracted from multiple stages (supplementary table S8) using the TRIzol plus RNA purification kit (Thermo Fisher Scientific).

### Genome and Transcriptome Sequencing

We performed short-read genome sequencing of 21 strains using Illumina HiSeq 2500/3000 sequencing (10× to 60× coverage, 100/150 bp paired-end; supplementary table S16). The library preparation and short-read sequencing were performed by BGI Japan (Kobe, Japan) or K. K. DNAFORM (Yokohama, Japan). The short-read data of the NIES-68 strain were previously published (accession number: DRR097270, Kanda et al. 2017). For the NIES-53 and NIES-54 strains, the library preparation and long-read sequencing of PromethION were performed by GeneBay (Yokohoma, Japan). The PacBio Sequel sequencing was performed for NIES-4552, NIES-4550, NIES-64, and NIES-65. In addition, 10X genomics linked reads of NIES-64 and NIES-65 were obtained from Macrogen (Kyoto, Japan). We obtained short-read paired-end transcriptome sequences of the six strains using MGI DNBSEQ-G400, MGI MGISEQ-2000, or Illumina HiSeq 2500 (supplementary table S8). The library preparation for transcriptome sequencing runs was performed by GENEWIZ Japan (Tokyo, Japan).

### Genome Size Estimation on *k*-mer Analysis

We performed a *k*-mer analysis on short-read sequences to estimate the genome size. We first removed reads with a Phred quality score ˂ 20 and reads length ˂ 50 bases using SolexaQA++ v3.1.7.1 (Cox et al. 2010). *k*-mer frequency distribution with *k* = 25 was calculated using JELLYFISH v2.2.4 with the - h 10000000 option for each strain (Marçais and Kingsford 2011). The *k*-mer occurrences due to error sequences were defined as smaller than the first local minimal value (*Xa*). The first peak of the *k*-mer occurrences was *Xb* (supplementary fig. S12). Therefore, we first estimated the genome size by calculating the total number of all *k*-mer from *Xa* to 20 × *Xb* divided by *Xb.* However, this method should be an underestimate because it excludes high copy (˃20) sequences.

Therefore, we also calculated the genome size without the upper limit of *k*-mer (supplementary table S3). Since the high copy sequences should include those from organellar genomes, we excluded them in the following way. We first estimated organellar genome size by *de novo* assembly of the organellar genomes from short-read sequences using GetOrganelle with the -k 21,25,55,85,115 -F embplant_mt,embplant_pt option (Jin et al. 2020). Mitochondrial and plastid genome sizes were estimated to be 98,950–193,473 and 139,599–158,119 bp, respectively. We then calculated the coverage ratio of organellar and nuclear genomes. We identified two peaks in high *k*-mer (˃500) indicating reads from mitochondrial (*Xm*) and plastid (*Xp*) genomes and divided them by *Xb*.

Using the genome of the NIES-4550 strain, the relative size of *Xm* and *Xp* (*Xm* ˂ *Xp*) was confirmed based on the following mapping. First, we concatenated the *de novo* assembled full genome and the assembled organellar genomes and mapped short-read sequences. Second, we calculated the depth of each position using the samtools depth with the -a -d 0 option (Li et al. 2009). Third, as the *de novo* assembled full genome also included contigs from organellar genomes, we searched the assembled organellar genomes against the *de novo* assembled full genome using BLASTN v2.0.15 with default parameters (Altschul et al. 1990) and identified the organellar contigs as the contigs with the number of identical matches ˃ 30,000 and the percentage of identical matches ˃ 99%. We then defined the coverage of organellar genomes as the sum of the average depth of these contigs and assembled organellar genomes. The coverage of mitochondrial and plastid genomes was 17,639.6 and 77,398.5, respectively, suggesting that the coverage of mitochondria is smaller than that of plastid.

### Investigation of the Inheritance of Genome Size Variation using Flow Cytometry

To establish the F_1_ hybrids between heterothallic strains, NIES-64 and NIES-65 strains were mixed and sexual reproduction was induced in an MI medium (1,000 cells each/200 μL) in 48-well plates. First, formed zygospores were cultured at 21°C under 24 h continuous light for three weeks. Next, we left zygospores at 21°C under dark conditions for a month to dry out fully. We then added 300 μL of a CA medium to induce germination of the zygospores. Following Tsuchikane et al. (2018b), vegetative cells from the germinated progenies were isolated and cultured in a conditioned CA medium, a sterilized CA medium in which cells were cultured for two weeks. For subsequent analysis, we used F_1_ lines that underwent 2‒5 times of subculturing every 10 days to two months, which would roughly correspond to the order of 100 generations of vegetative reproduction. Note that this subculturing period was relatively short compared to those of the parental strains maintained at the NIES stock center, which were subcultured every 10 days to four months since 1974 (Kasai et al. 2004).

To understand how genome size variation between strains is inherited, the genome size of the F_1_ lines was estimated by flow cytometry. First, 50,000 cells were washed with a working solution (0.4 M sorbitol and 30 mM 2(N-Morpholino) ethane sulfonic acid), then incubated using a 100 μL enzymatic mixture of 0.5% Cellulase Onozuka R-10 and 0.5% Macerozyme R-10 dissolved in the working solution at 29°C for 30 min. Next, samples were centrifuged at 14,000 rpm for five minutes, and the supernatant was decanted. To isolate nuclei, we added 500 μL of Cystain PI Absolute P nuclei extraction buffer (Sysmex) and filtered through a 20 μm filter (2-7209-01; AS ONE CORPORATION, Osaka, Japan).

We used *Lycopersicon esculentum* ‘Stupicke’ as a control (Dolezel et al. 1992). Young leaf (∼1 mg) was mixed with 150 μL of Cystain PI Absolute P nuclei extraction buffer and chopped with a razor for three minutes, then the sample was filtered through a 20 μm filter. Next, we added 1,200 μL Cystain PI Absolute P staining solution (Sysmex), which contained 12 μg RNase A to digest RNA, to the filtrate of *C. psl.* complex nuclei and control nuclei, and left it for 3 h at 4°C.

Fluorescence was measured using a BD Accuri C6 flow cytometer (BD Biosciences) equipped with a 488 nm laser. Measurement was performed at least three times per strain. We obtained fluorescence intensity at FL2 = 585 ± 20 nm and signal width as the size of the particles. We analyzed the output FCS files using the R package flowCore (Hahne et al. 2009). We excluded particles with a width ˂ 40 or ˃ 80 and detected peaks as nuclear DNA content. Because the peak representing 2C was detected more prominently than 1C in all strains, we estimated the 1C genome size from the second peak. We confirmed that these estimates were strongly correlated with the observed values for the first peak, 1C (*R*² = 0.912; supplementary fig. S13). The genome size of each sample was estimated based on that of *L. esculentum* ‘Stupicke’ (1C = 958 Mb, Dolezel et al. 1992; supplementary table S4). Strains measured at least three times were utilized to compare the variance within and among strains using the R package heritability (Kruijer 2016), and finally, the unimodality test was conducted with modetest in the R package multimode (Ameijeiras-Alonso et al. 2021; fig. 3). Additionally, strains with less than three measurements were included when we compared the genome size of F_1_ lines with the minimum number of subculturing (three times, corresponding to about 50–200 generations of vegetative reproduction), which were isolated on 23-Jun-2021 and measured on 6-Jan-2022 (supplementary fig. S2 and table S4).

### *De novo* Assembly

NIES-4552 was assembled with Canu 1.5 (Koren et al. 2017) with genomeSize=1g and -pacbio-raw options. Around 4.5 million subreads with N50 of 17 kb totaling 44.8 Gb were used. We used ntEdit v1.3.5 (Warren et al. 2019) to polish the assembly using DNA short-read data from the NIES-4552. NIES-64, NIES-65, and NIES-4550 were assembled with Canu 1.8 (genomeSize=1.2 g) and the error were corrected with Arrow (PacificBiosciences). For NIES-64, 5.2 million reads with N50 of 30 kb totaling 107 Gb; for NIES-65, 8.9 million reads with N50 of 30 kb totaling 110 Gb; for NIES-4550, 5.7 million reads with N50 of 27.9 kb totaling 107 Gb of raw reads were used. For NIES-64 and NIES-65, the arrow-corrected contigs were scaffolded with 10x genomics linked read data using ARCS (Yeo et al. 2018) after mapping with LongRanger-2.2.2 (10x Software). The reads are mapped again and used for error correction using Pilon for indel errors (Walker et al. 2014). For NIES-64, after inspecting the relationship between the mapped tag number and the reference length, contigs with more than 1 tag per 100 bp reference were retained as authentic contigs (supplementary fig. S14). The rest may represent bacterial or other microbial contamination in the culture. NIES-4550 assembly was error corrected for indel errors with short-read mapping by BWA MEM 0.7.15-r1140 (Li and Durbin 2009) using Pilon. NIES-53 and NIES-54 were assembled with Platanus-allee v2.2.2 with short read and PromethION data (Kajitani et al. 2019). Here, the short reads were trimmed with Trimmomatic 0.39 (Bolger et al. 2014), specifying ILLUMINACLIP:TruSeq3-PE.fa:2:30:10 LEADING:3 TRAILING:3 SLIDINGWINDOW:4:15 MINLEN:36 before to assembly and only paired reads were used. A total of 55.4 and 59.4 Gb were retained for NIES-53 and NIES-54 after trimming, respectively. Nanopore long reads were 1 million reads with N50 of 25.6 kb totaling 14.8 Gb for NIES-53 and 0.76 million reads with N50 of 31.6 kb totaling 12.4 Gb for NIES-54.

For all six assembled genomes, we mapped their paired-end short reads to the assembled genomes of themselves using BWA MEM v.0.7.17 with default parameters (Li and Durbin 2009) and obtained depth information using DepthOfCoverage of GATK v.3.8-1-0-gf15c1c3ef with the default option (McKenna et al. 2010). We then removed the contigs where the percentage identity of nonzero coverage was smaller than 0.001.

### Observation of Nuclei

Cells cultured for approximately 10 days were collected by centrifugation and fixed with a solution of ethanol and acetic acid (3:1) for 20 minutes. The fixed cells were then washed with 1x Phosphate Buffered Saline (PBS) and stained with 1µg/mL DAPI in 1x PBS, followed by observation under a UV fluorescence microscope. The area of nuclei in focus was measured using ImageJ (Schneider et al. 2012).

### Mapping of DNA Short Reads

We obtained a set of SNPs of all strains to construct a phylogenetic tree. Paired-end short reads filtered by SolexaQA++ v3.1.7.1 (Cox et al. 2010) were mapped to *de novo* assembled sequences of the NIES-4552 strain as a reference using BWA MEM v.0.7.17 with default parameters (Li and Durbin 2009). To preprocess generated SAM files for GATK v4.0.4.0 (McKenna et al. 2010), we converted SAM files to BAM files using SortSam, AddOrReplaceReadGroups, and MarkDuplicates on Picard v.2.27.1 (http://broadinstitute.github.io/picard/). VCF files were created using HaplotypeCaller of GATK v4.0.4.0. In the VCF file created by mapping to the NIES-4552 strain, information of 79,377 SNPs was obtained from all strains using the package VCFR in R (Knaus and Grünwald 2017). Heterozygous SNPs were excluded. We applied the ASC_GTRGAMMA model using RAxML v.8.2.12 with the -f a -m ASC_GTRGAMMA --asc-corr lewis -p 2 -# 100 -x 12345 option (Stamatakis 2014).

Using the same method as presented above, we also mapped paired-end short reads of NIES-58, NIES-59, NIES-64, and NIES-65 to *de novo* assembled sequences of the NIES-4550 strain. We obtained depth information using DepthOfCoverage of GATK v.3.8-1-0-gf15c1c3ef with the default option. To understand the differences in depth of each strain by gene, we calculated the average depth of each gene and standardized the coverage by the average depth among the strains.

To identify CNV between NIES-64 and NIES-65 from the DNA depth log-ratio, we mapped paired-end short reads of NIES-64 and NIES-65 to *de novo* assembled sequences of the NIES-65 using HISAT2 version 2.1.0 with the --no-spliced-alignment option (Kim et al. 2015). We counted the reads mapped on the exon regions using FeatureCounts v2.0.1 with the -p -t exon option (Liao et al. 2014). The read counts were scaled based on the DNA depth per million (DPM). DPM was calculated by the following way: we first divided the read counts by the length of each gene in kilobases (read per kilobase; RPK). Then RPK was divided by the sum of RPK for all genes in a million bases. We summed up the DPM for each OG. The DNA depth log-ratio of each OG was defined as the difference in log_2_-transformed DPM of each OG (NIES-65 – NIES-64). We changed the criteria of the DNA depth log-ratio for CNV depending on the total DNA depth, i.e., the sum of the normalized log_2_-transformed DPM of each OG of NIES-64 and NIES-65. This is because the values of nonzero peaks, indicating CNV, differed depending on the total DNA depth (fig. 6b, supplementary fig. S9). This pattern would be expected since the DNA depth ratio should differ depending on the total copy number of two strains. For example, when the total copy number of an OG is three copies (total DNA depth: ∼75), the expected DNA depth ratio should be 1:2 or 2:1, corresponding to the DNA depth log-ratio as –1 or 1. When the total copy number of an OG is five (total DNA depth: ∼125), the expected DNA depth ratio should be 2:3, 3:2, 1:4, or 4:1, corresponding to the DNA depth log-ratio as –0.58, 0.58, –2, or 2. Note that the mean DPM of a single copy gene was 24.64. We considered that a gene has CNV if |DNA depth log-ratio| ˃ 0.5 and a total DNA depth ˂ 90, or if |DNA depth log-ratio| ˃ 0.36 and a total DNA depth ≥ 90.

### Gene Prediction

First, we constructed a repeat model for each strain using RepeatModeler Version open-1.0.11 (Smit and Hubley 2015) with the National Center for Biotechnology Information (NCBI) engine and identified repeat sequences using RepeatMasker version open-4.0.7 (Smit et al. 2015; supplementary table S5).

Then, genome annotation was performed independently for each strain using EVidenceModeler v.1.1.1 (EVM; Haas et al. 2008). EVM created consensus gene structures from Augustus v.3.3 (Keller et al. 2011) with weight 1 and Program to Assemble Spliced Alignments (PASA) v2.4.1 with weight 20 and 10 in the NIES-4552 genome and the others, respectively (Haas et al. 2003). We first performed training for Augustus using BUSCO v4.0.2 with the --long option based on viridiplantae_odb10 (Simão et al. 2015). We then predicted gene locus using three hint files. The first hint file was generated from the RNA-seq data. After creating a database file of masked genome sequences using Bowtie2 v2.2.6 (Langmead and Salzberg 2012), we mapped raw reads from RNA-seq to the sequences using TopHat v.2.1.0 (Kim et al. 2013). The output was converted to the appropriate hint files created following Bioinformatics Greifswald Software Wiki and Forum (https://bioinf.uni-greifswald.de/bioinf/wiki/pmwiki.php?n=IncorporatingRNAseq.Tophat). The second hint file was based on the amino acid sequences of other species; *Arabidopsis thaliana* (TAIR10_pep_20110103_representative_gene_model_updated), *Physcomitrella patens* (P.patens.V6_filtered_cosmoss_proteins.fas), *Chara braunii* (Nishiyama et al. 2018), *Spirogloea muscicola* and *Mesotaenium endlicherianum* (Cheng et al. 2019). Then, the amino acid sequences were mapped using Exonerate (Slater and Birney 2005) v2.4.0 and converted to a hint file using exonerate2hints.pl in Augustus. The third hint file was made from RepeatMasker output following Greifswald Software Wiki and Forum (https://bioinf.uni-greifswald.de/bioinf/wiki/pmwiki.php?n=Augustus.IncorporateRepeats). On PASA, we built a comprehensive transcriptome database using two transcriptome assembly datasets. The first dataset comprised genome-guided transcriptome assembly data produced using Trinity v.2.1.1 with the -- genome_guided_max_intron 10000 option (Grabherr et al. 2011) based on the RNA-seq data mapped to the masked genome sequences using HISAT2 version 2.1.0 (Kim et al. 2015). The second dataset comprised *de novo* assembled transcriptome data produced using Trinity. We used the BEDTools complement v2.26.0 (Quinlan and Hall 2010) to obtain lists of the length of intron and intergenic regions.

### Dot Plot of Self-alignment

We performed self-alignment of four *de novo* assembled genomes, NIES-4552, NIES-64, NIES-65 and NIES-4550, using D-GENIES with the “many repeats” option (access date: Jan. 2023; Cabanettes and Kloop 2018). From the output files, we deleted the results between contigs that are identical, and plotted the alignment using D-GENIES (access date: Jan. 2023).

### Copy Number Count and Topology Analysis for Each Ortholog Group

OGs were identified using amino acid sequences of annotated genes for the six strains. First, we used SonicParanoid with the -I 3 option (fine-grained clustering; Cosentino and Iwasaki 2019) and then identified 29,956 OGs. The output files were used to count the copy number for each OG. Next, we calculated the mean copy number and proportion of genes with CNV per contig using genes with copy numbers ˂ 10 (fig. 4).

We compared the distributions of synonymous substitution rates (*K*_s_) for all combinations of genes among strains and within each OG. *K*_s_ of all combinations of genes within each strain were calculated using wgd v1.1, showing peaks at approximately *K*_s_ = 0.35 (Zwaenepoel and Van de Peer 2019; supplementary fig. S15a). Maximum *K*_s_ values for all combinations within each OG were calculated using the R package seqinR (Charif and Lobry 2007). The position of the first peak of the within-OG distribution mostly corresponded to the peak of the pairwise genome-wide *K*_s_ distribution, suggesting that the OGs defined by the SonicParanoid software generally clustered orthologous genes (supplementary fig. S15b). Therefore, we only used OGs with the maximum *K*_s_ values ˂ 1 in subsequent analyses to eliminate OGs containing paralogs that diverged too early.

To identify the origin of duplicated genes, we constructed phylogenetic trees of coding sequences (CDS) from the same OG where zero to four copies of genes were found in all strains. First, we aligned CDS using MAFFT v7.305b with default parameters (Katoh and Standley 2013). We then excluded all sites with gaps from the aligned sequences and constructed phylogenetic trees using IQ-TREE version 1.0 with the -m TIM3+F+I -B 1000 -keep-ident option (Nguyen et al. 2015). Finally, we evaluated the bootstrap values and the topologies of phylogenetic trees using check_monophyly in the package ete3 (Huerta-Cepas et al. 2016) in Python 3.7. To identify the origin of duplicate genes in the NIES-4550 strain, we focused on the Ogs where NIES-4550 had two copies and the five other strains had a single copy, and treated the NIES-4552 strain as an outgroup (supplementary table S6). When we identified the origin of duplicate genes in five other strains, we removed the leaf nodes from NIES-4550 from the tree and focused on the Ogs where all five strains had two to four copies (supplementary table S7).

### Amplification and Sequencing the Putative Recombination Breakpoints

To experimentally validate the putative recombination breakpoints between regions with different presumptive origins, we amplified the genomic regions encompassing the breakpoints by PCR and performed Sanger sequencing (supplementary table S17). We chose regions in which genes with different presumptive origins were adjacent to each other within 10 kb (supplementary table S17). PCR was performed using TaKaRa PrimeSTAR GXL DNA Polymerase following the manufacturer’s protocol (Takara Biotechnology, Japan). Amplified DNA fragments were purified by illustra ExoProStar (GE Healthcare Life Sciences). PCR-amplified fragments with the sequencing primers were shipped to Eurofins Genomics K.K. (Tokyo, Japan) for Sanger sequencing (supplementary table S17).

### Gene Expression Analysis

To investigate the relationship between copy number and gene expression in the strains NIES-64 and NIES-65, we obtained RNA-seq data with three replicates from cells during five stages: during vegetative reproduction (morning), during vegetative reproduction (night), under a nitrogen-depleted condition, and in conditioned MI medium for 8 h and 24 h (fig. 1; see supplementary table S8 for details). The conditioned an MI medium was MI medium in which a mixture of NIES-64 and NIES-65 had been cultured for 48 h. In addition, we used cells that had been vegetatively grown for ten days for all stages.

To preprocess paired-end raw reads, we applied fastp (Chen et al. 2018) with the -trim_poly_x option. Next, filtered reads were mapped to *de novo* assembled genome sequences of NIES-65 using HISAT2 version 2.1.0 with the default option (Kim et al. 2015). We then counted reads mapped on exons using FeatureCounts v2.0.1 with the -p -t exon option (Liao et al. 2014). To compare expression levels between stages, we produced TMM-normalized data using the package edgeR in R (Robinson et al. 2010). Next, the read counts were scaled based on Transcript per million (TPM). TPM was calculated by the following way: we first divided the read counts by the length of each gene in kilobases (read per kilobase; RPK). Then RPK was divided by the sum of RPK for all genes in a million bases. Finally, we summed up the TPM-scaled read counts mapped to each OG, and then defined OGs with TPM ˃ 5 in all five stages as genes with constitutive expression, and genes with TPM ˂ 0.1 in at least one stage and TPM ˃ 5 in at least one another stage as genes with conditional expression (Table S15).

We determined DEGs with FDR ˂ 0.05 using the package edgeR in R (Robinson et al. 2010). We performed Fisher’s exact tests to examine if DEGs were overrepresented in genes with CNV. To identify dosage-compensated genes, we performed the two following tests. First, to test whether the expression differences were smaller than the expected copy number differences, we performed one-sided *t*-tests with DNA depth ratio for the log_2_-transformed expression data of the OGs with CNV. We regarded that OGs with FDR ˂ 0.05 had significantly smaller expression level differences than the expected gene dosage differences. Second, to test whether the OGs showed the inverse effect, i.e., a significantly higher expression in the low copy number strain than in the high copy number strain, we performed one-sided *t*-tests for the log_2_-transformed expression data. We identified OGs with FDR ˃ 0.1 as OGs without a negative dosage response.

### GO Analysis

GO term enrichment analysis was performed to test if any GO is enriched in the various gene sets of interest. To determine the GO terms of the predicted genes, we searched the amino acid sequences of genes against the NCBI nonredundant protein database using DIAMOND BLASTP v2.0.15 with default parameters (Buchfink et al. 2021) and the UniProtKB/Swiss-Prot database using BLASTP v2.6.0+ with the -evalue 1e-5 -max_target_seqs 1 option (Altschul et al. 1990). We assigned the 7,666 GO annotations of the 22,809 genes on NIES-65. Then, GO terms enrichment analysis was conducted using topGO (Alexa et al. 2006) and GO.db v.3.12.1 (Carlson et al. 2019) in R. We used Fisher’s exact test with the ‘weight01’ algorithm in the R package topGO to reduce redundancies for the statistical significance measurements with a significance level of ˂0.05 (Alexa et al. 2006).

## Acknowledgments

This work was supported by JSPS KAKENHI (Grant Numbers 15K18583, 17K15165, and 16H06279 [PAGS] to TT, 20J20877 to YWK, 19K06827 to YT, and 16H04836 to HS) and by NODAI Genome Research Center. Computations were partially performed on the NIG supercomputer at ROIS National Institute of Genetics. We are grateful to Dr. Hiroyuki Koga for flow cytometry analysis.

## Data Availability

Transcriptomic sequence data from the NIES-4550 strain have been deposited in the DNA Data Bank of Japan (DDBJ) under DRA003074. In addition, other raw genomic and transcriptomic sequence data have been deposited in the European Nucleotide Archive (ENA) under BioProject PRJEB57781.

