## Supplementary Note and Figures for "Extensive Copy Number Variation Explains Genome Size Variation in the Unicellular Zygnematophycean Alga, *Closterium peracerosum-strigosum-littorale* Complex"

### Supplementary Materials

Yawako W. Kawaguchi[1,2*], Yuki Tsuchikane[1,3], Keisuke Tanaka[4], Teruaki Taji[5], Yutaka Suzuki[6], Atsushi Toyoda[7], Motomi Ito[8], Yasuyuki Watano[9], Tomoaki Nishiyama[10], Hiroyuki Sekimoto[3] & Takashi Tsuchimatsu[1*]

[1]Department of Biological Sciences, Graduate School of Science, The University of Tokyo, 7-3-1 Hongo, Bunkyo-ku, 113-0033 Tokyo, Japan

[2]Graduate School of Science and Engineering, Chiba University, 1-33 Yayoi-cho, 263-8522 Chiba, Japan

[3]Department of Chemical and Biological Sciences, Faculty of Science, Japan Women's University, 2-8-1 Mejirodai, Bunkyo-ku, 112-8681 Tokyo, Japan

[4] NODAI Genome Research Center, Tokyo University of Agriculture, 1-1-1 Sakuragaoka, Setagaya-ku, 156-8502 Tokyo, Japan

[5]Department of Bioscience, Tokyo University of Agriculture, 1-1-1 Sakuragaoka, Setagaya-ku, 156-8502 Tokyo, Japan

[6]Graduate School of Frontier Sciences, University of Tokyo, 5-1-5 Kashiwanoha, Kashiwa, 277-8568 Chiba, Japan

[7]Advanced Genomics Center, National Institute of Genetics, Mishima, 411-8540 Shizuoka, Japan

[8]Graduate School of Arts and Sciences, The University of Tokyo, 3-8-1 Komaba, Meguro-ku, 153-8902 Tokyo, Japan

[9]Graduate School of Science, Chiba University, 1-33 Yayoi-cho, 263-8522 Chiba, Japan

[10]Research Center for Experimental Modeling of Human Disease, Kanazawa University, Takaramachi 13-1, Kanazawa, 920-0934 Ishikawa, Japan

**Supplementary Note: The NIES-4550 Genome Contains Sequences from the Three Different Origins**

We found that the NIES-4550 genome contains genes from at least two different origins. This is because the coverage depth of reads from groups II-A and II-B showed a complementary pattern and because we identified the genes of the NIES-4550 genome forming a single clade with genes of the mating group II-A or II-B (fig. 5a, supplementary table S6, see main text for details). To identify whether each gene of the NIES-4550 strain originated from group II-A or II-B, we calculated the genetic distance between the NIES-4550 strain, the group II-A, and the group II-B based on the phylogenetic tree for each OG. Genetic distance was obtained as the maximum likelihood distance from the software IQ-TREE version 1.0 with the -m TIM3+F+I -B 1000 -keep-ident options (Nguyen et al. 2015). We used only OGs with aligned sequences > 500 bp in length. The genetic distance of each combination between the sequences from the NIES-4550 strain and the group II-A was denoted by $D_{NAn}$ ($1 \leq n \leq$ the copy number of genes from the group II-A), and the average genetic distance of all these combinations was denoted by $D_{NA}$. We also calculated $D_{NBn}$, $D_{NB}$, and $D_{AB}$, respectively, the distances of each combination between the NIES-4550 strain and the group II-B, the average distance between the NIES-4550 and the group II-B and the average distance between groups II-A and II-B (supplementary fig. S16a). We then calculated $D$, the difference between $D_{NA}$ and $D_{NB}$ normalized by $D_{AB}$, to evaluate whether a gene from the NIES-4550 genome was closer to group II-A or II-B (supplementary fig. S16b). If a gene from NIES-4550 is closely related to the genes from group II-A or II-B, a value of $D$ should be smaller or larger than zero, respectively (supplementary fig. S16b left and right). By contrast, if a sequence from NIES-4550 is highly distant from groups II-A and II-B, $D$ should be close to zero (supplementary fig. S16b center). We detected a trimodal distribution of $D$ and identified 4,727 genes with $D$ close to zero ($-0.4 < D < 0.4$, supplementary fig. S17a). We defined these genes as “unclassified genes.”

There are two possible explanations for the unclassified genes: (1) mis-assembly and/or recombination between genes from groups II-A and II-B; and (2) the presence of genomic regions from the third major ancestor other than groups II-A and II-B. To test these two possibilities, we calculated the relative number of mutations unique to NIES-4550 ($U$ = (the average of $D_{NAn}$ and $D_{NBn}$)/$D_{AB}$, supplementary figs. S16c and S17b and c). If a sequence does not harbor any mutation unique to NIES-4550, $U$ is 0.5. If the unclassified genes originated from mis-assembly and/or recombination (1), the number of unique mutations of unclassified genes would be comparable to that of classified genes. If the unclassified genes originated from the third major ancestor (2), the number of unique mutations of unclassified genes would be larger than that of the classified genes. We found that the mode of the distribution of the unclassified genes was at 0.76, which is larger than that of the classified genes at 0.52 (supplementary fig. S17c), suggesting that unclassified genes originated from the third major ancestor.

We defined the genes with $D < -0.6$ and $U < 1$ as genes from the group II-A (2,440 genes), the ones with $D > 0.6$ and $U < 1$ as those from the group II-B (2,717 genes), and the ones with $D > -0.4$, $D < 0.4$ and $U < 1.5$ as those from the third major ancestor (4,250 genes). We then investigated the distribution of these genes with different origins along the contigs; we found that these genes with different origins were sometimes found in a single contig and adjacent to each other, suggesting recombination between genomic fragments of different origins (supplementary fig. S7a and b). To experimentally test whether these genes with different origins are physically adjacent, we performed a PCR that amplifies genomic fragments encompassing putative recombination breakpoints. We confirmed the amplification of these fragments in two independent putative recombination breakpoints and their nucleotide sequences using Sanger sequencing (supplementary figs. S7c and S8). In summary, these results suggest that the heterogeneity of the NIES-4550 genome reflects its hybrid origin with further subsequent rearrangements including recombination between genomic segments with different origins.

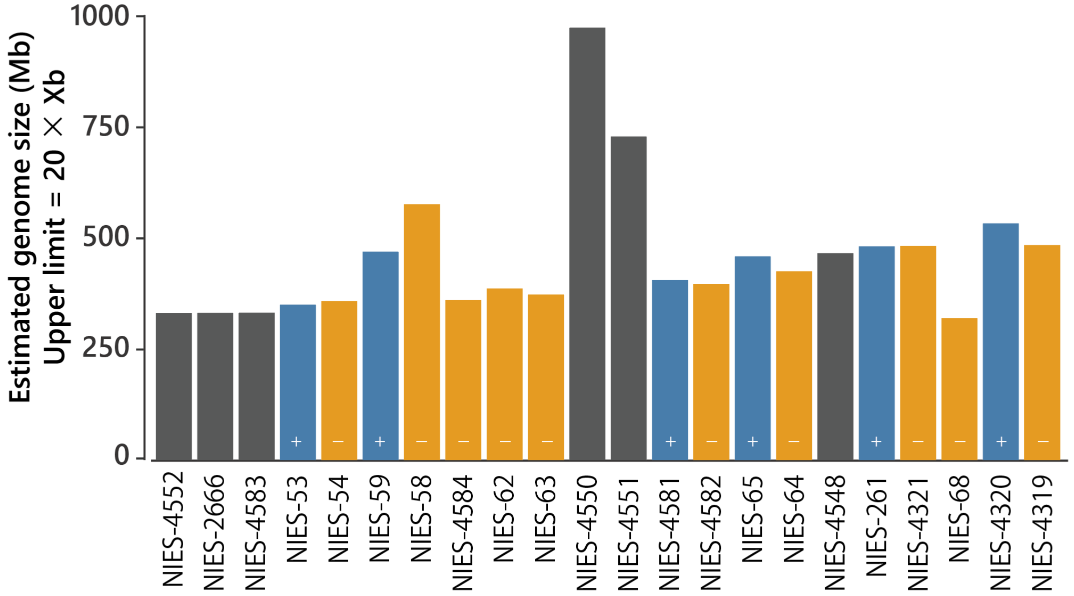


**Fig. S1. Estimated genome size variation by *k*-mer analysis with upper limit = 20 × *Xb* in the 22 strains of the *Closterium peracerosum-strigosum-littorale* complex.**

*Xb* indicates the first peak of the *k*-mer occurrences (see Materials and Methods and supplementary fig. S12 for details). Black, blue, and orange bars indicate the estimated genome size of homothallic, heterothallic (mt$^+$), and heterothallic (mt$^-$) strains, respectively.

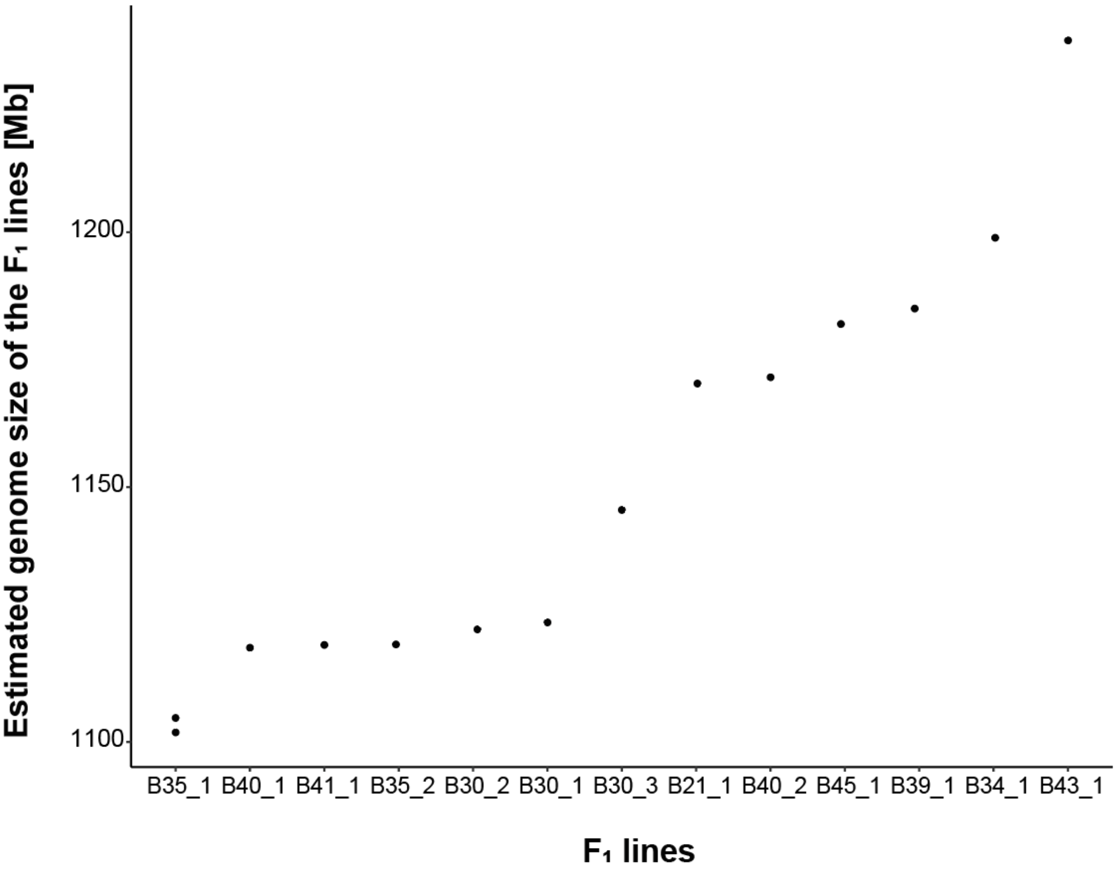


**Fig. S2. Genome size variation by flow cytometry among $F_1$ lines with the minimum number of subculturing.**

Genome size of each $F_1$ line subcultured three times (corresponding to about 50–200 generations of vegetative reproduction).

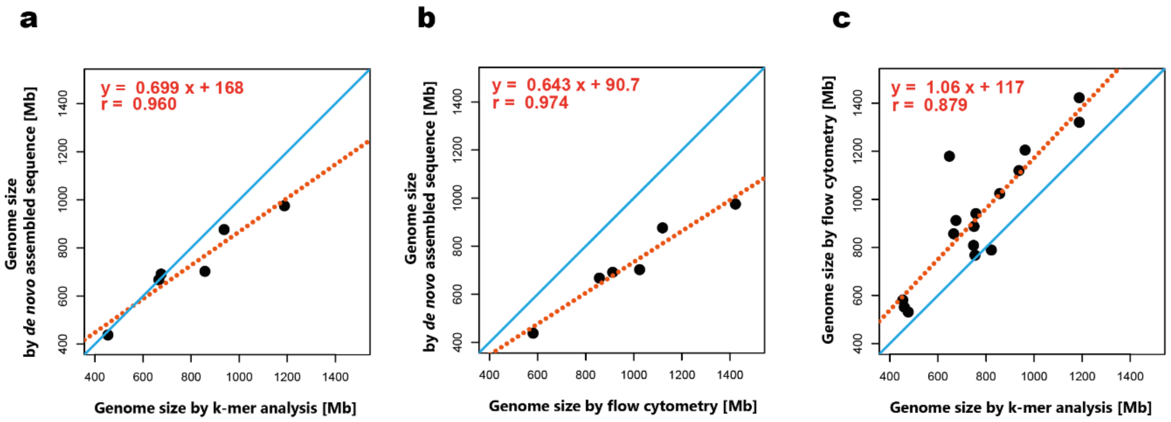


**Fig. S3. Genome size estimated by *de novo* assembled sequences, *k*-mer analysis and flow cytometry.**

(a-c) The estimated genome size based on *k*-mer analysis vs. *de novo* assembled sequences (a), flow cytometry vs. *de novo* assembled sequences (b), and flow cytometry vs. *k*-mer analysis (c). Light blue lines indicate $y = x$, while red dashed lines represent the simple regression. The estimated genome size by *k*-mer analysis is based on the calculation of the total number of all *k*-mer without the upper limit and the subtraction of organelle genome peaks (a and c).

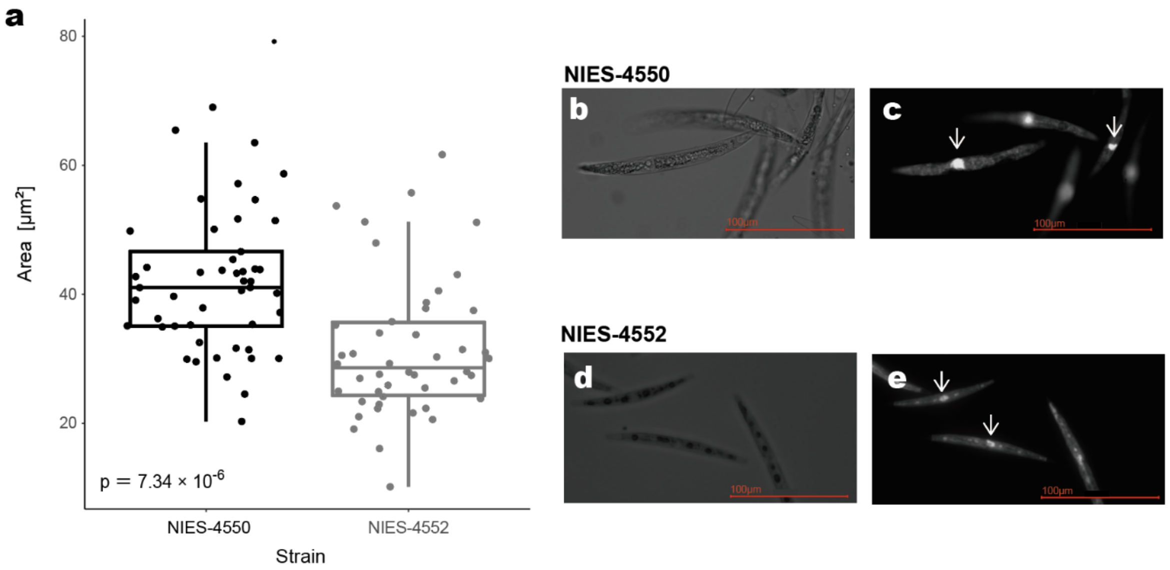


**Fig. S4. The nuclear size of the NIES-4550 and NIES-4552 strains.**

(a) The nuclear size of the NIES-4550 (estimated genome size: 975 Mb) was significantly larger in than that of the NIES-4552 (estimated genome size: 438 Mb). (b-e) Photographs of vegetative cells of the two strains under dark-field (b and d) and fluorescence (c and e) microscopy. (c, e) The area of nuclei in focus (indicated by arrows) was measured using ImageJ (Schneider et al. 2012).

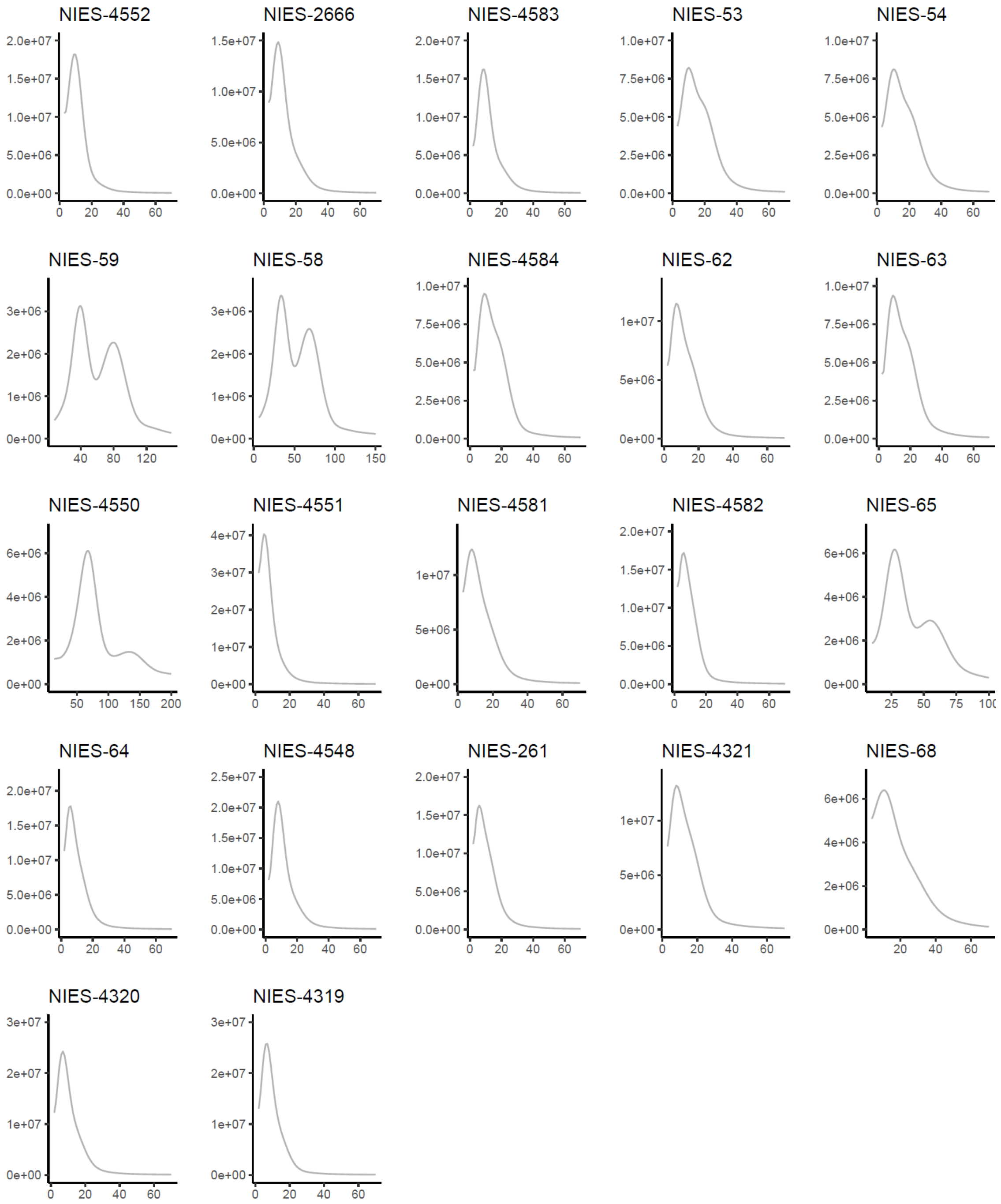


**Fig. S5. *k*-mer frequency distributions of the 22 strains with *k* = 25.**

Each panel shows *k*-mer frequency distributions trimmed the *k*-mer occurrence smaller than the first local minimal value.

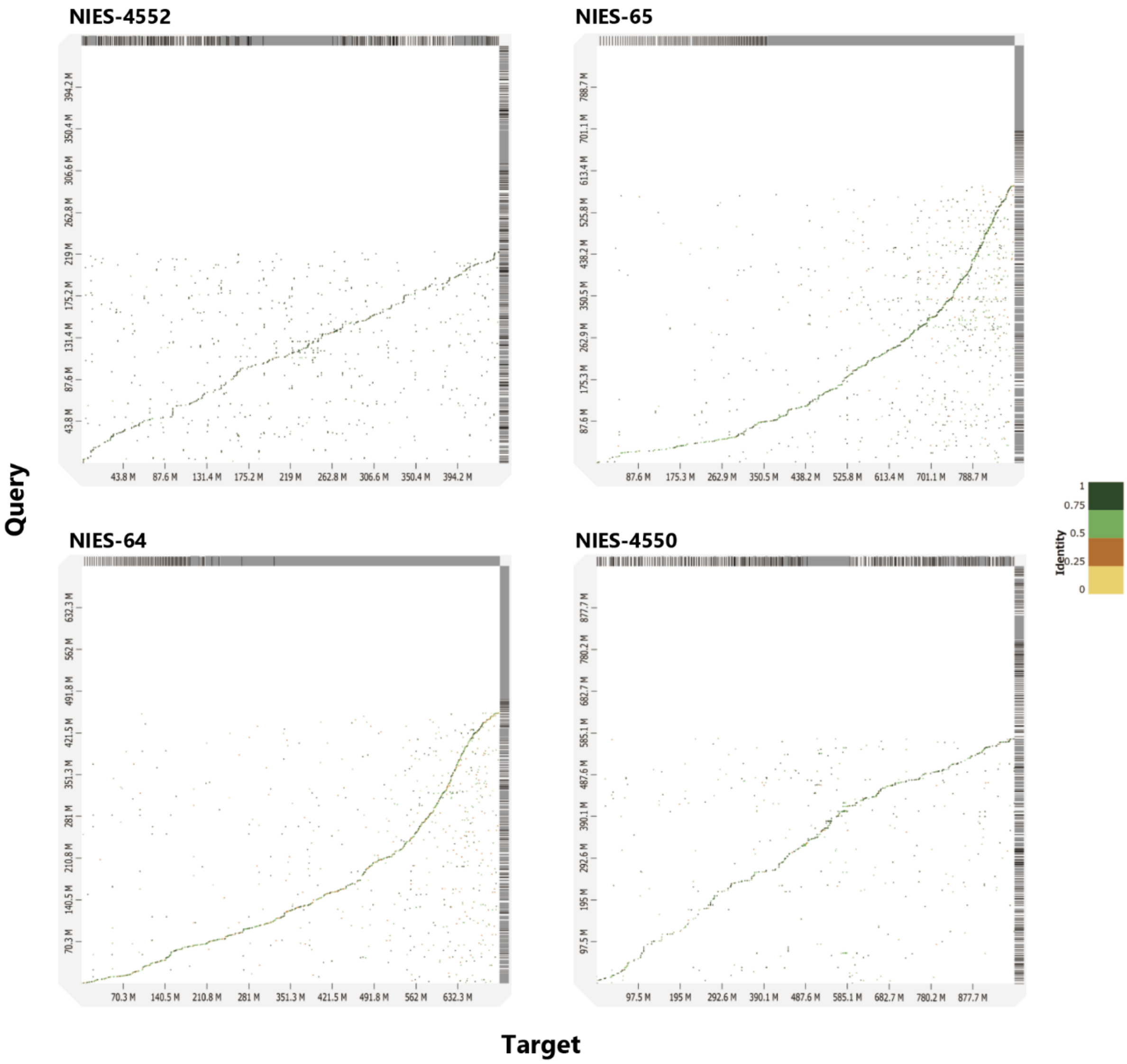


**Fig. S6. Dot plots of self-alignments.**

Each panel displays the results of self-alignments in each *de novo* assembled genome. Comparisons between identical contigs are not included. The aligned segments are represented as dots. The colors indicate similarity values that were binned in four groups (< 25, 25–50%, 50–75%, and > 75% similarity)

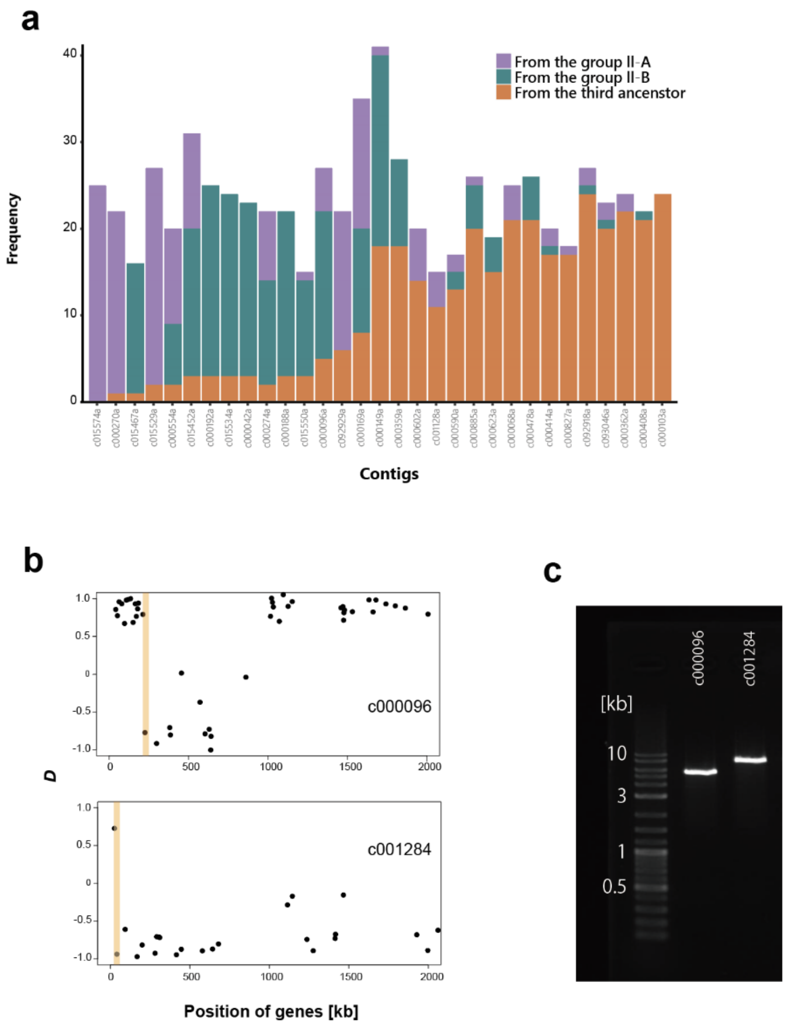


**Fig. S7. Recombination between genomic fragments of different origins in the NIES-4550 genome.**

(a) Classification of different origins for each contig. We show 31 representative contigs that have the largest number of classified genes. We defined the genes with $D < -0.6$ and $U < 1$ as genes from the group II-A (purple), the ones with $D > 0.6$ and $U < 1$ as those from the group II-B (green), and the ones with $D > -0.4$, $D < 0.4$ and $U < 1.5$ as those from a third major ancestor (orange). (b) $D$ statistics of each gene along two contigs, c000096 and c001284. Orange areas indicate the putative recombination borders between genomic fragments of different origins. (c) PCR amplifies the putative recombination borders in two contigs, c000096 and c001284, using a DNA ladder marker (DNA Ladder One Broad Range, Nacalai Tesque, Kyoto, Japan).

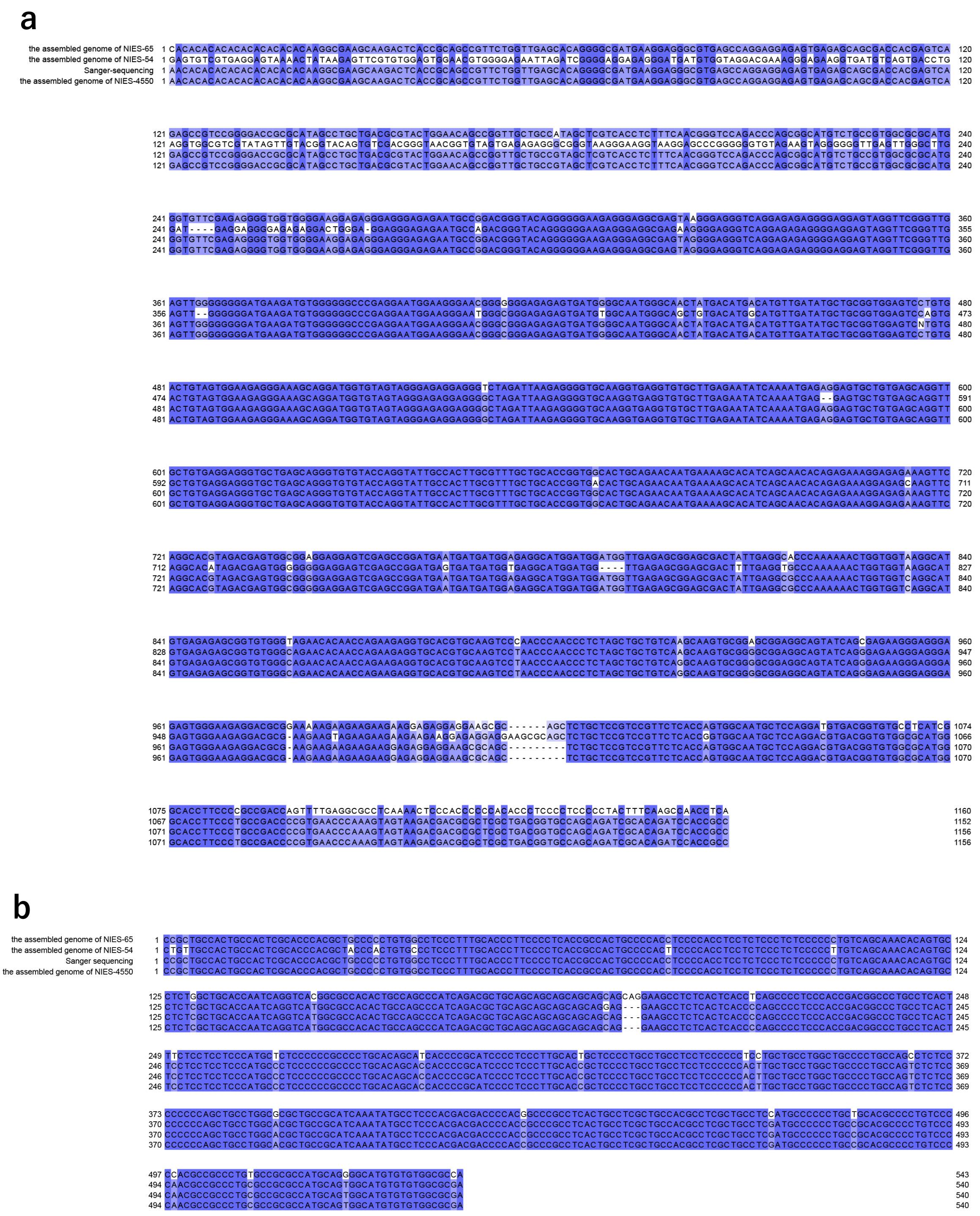
a
the assembled genome of NIES-65
the assembled genome of NIES-54
Sanger-sequencing
the assembled genome of NIES-4550
b
the assembled genome of NIES-65
the assembled genome of NIES-54
Sanger sequencing
the assembled genome of NIES-4550

**Fig. S8. The nucleotide sequences of two putative recombination breakpoints of the NIES-4550 genome.**

Regions from two contigs, c000096 (a) and c001284 (b) are shown. In both contigs, the assembled genome of NIES-4550 contains NIES-65-like and NIES-54-like sequences. The sequences of the assembled genome of NIES-4550 were confirmed by Sanger sequencing.

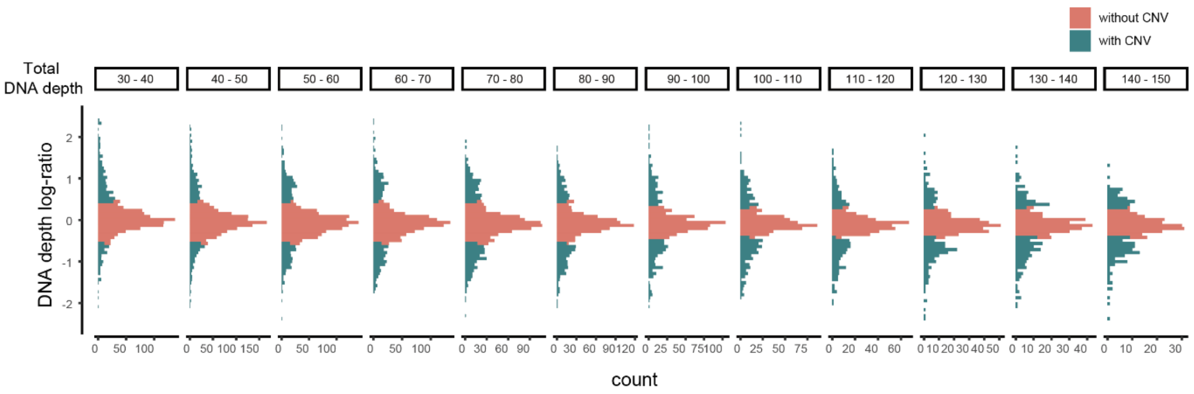


**Fig. S9. Distribution of DNA depth log-ratio between the NIES-64 and NIES-65 strains, depending on the total DNA depth.**

Each panel shows the distribution of the DNA depth log-ratio of OGs with the different total DNA depth. Green and red bars indicate OGs with and without copy number variation (CNV), respectively. We defined CNV as OGs with a total DNA depth < 90 and |DNA depth log-ratio| > 0.5, or with a total DNA depth ≥ 90 and |DNA depth log-ratio| > 0.36.

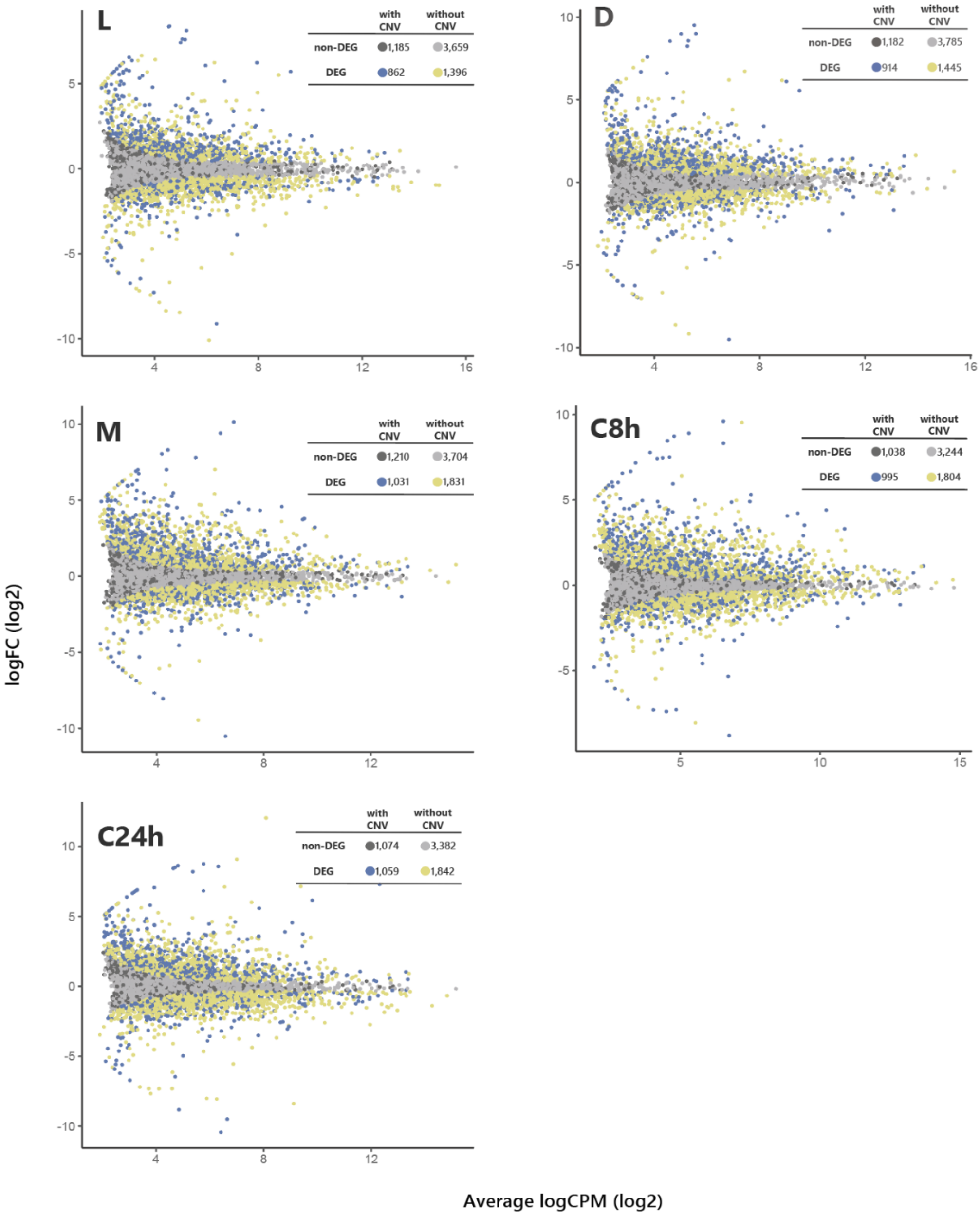


**Fig. S10. MA plots of gene expression difference between the NIES-64 and NIES-65 strains of each stage.**

Each panel shows results during each stage: during vegetative reproduction in light (L), during vegetative reproduction in dark (D), under a nitrogen-depleted condition (M), and in the mating-induced conditioned medium for 8 h (C8h) and 24 h (C24h). The number of genes and color in each category, differentially expressed genes (DEGs) or not, and with or without copy number variation (CNV), is shown in the upper right table of each panel. The genes with CNV were defined based on the DNA depth log-ratio.

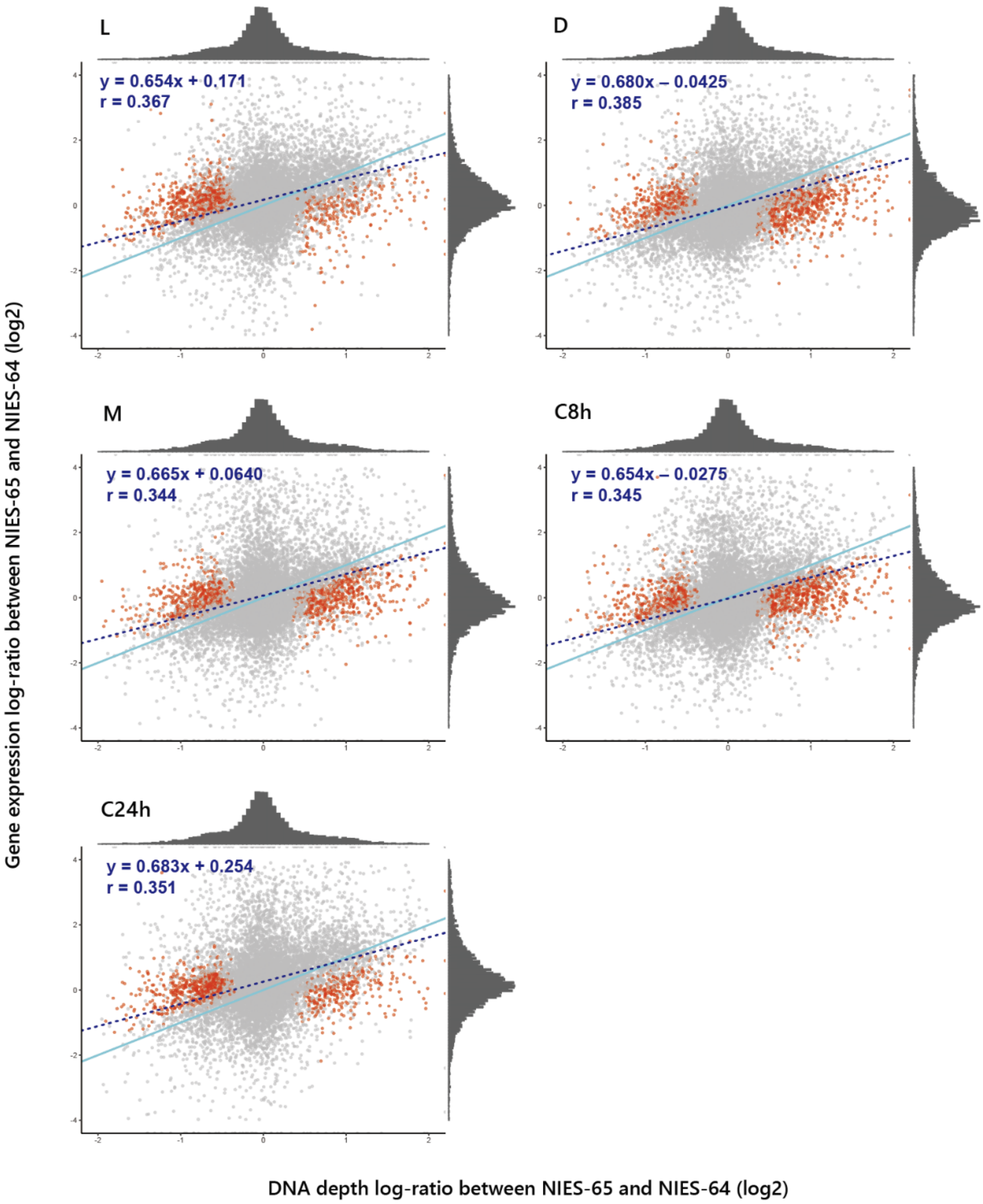


**Fig. S11. The expression level and gene dosage difference between each stage's NIES-64 and NIES-65 strains.**

Dosage-compensated genes are highlighted in red. A light blue line indicates the expected relationship representing proportionate increases in expression difference to dosage difference, while a blue dashed line represents the simple regression. Each column shows results during each stage; during vegetative reproduction in light (L), during vegetative reproduction in dark (D), under a nitrogen-depleted condition (M), and in the mating-induced conditioned medium for 8 h (C8h) and 24 h (C24h).

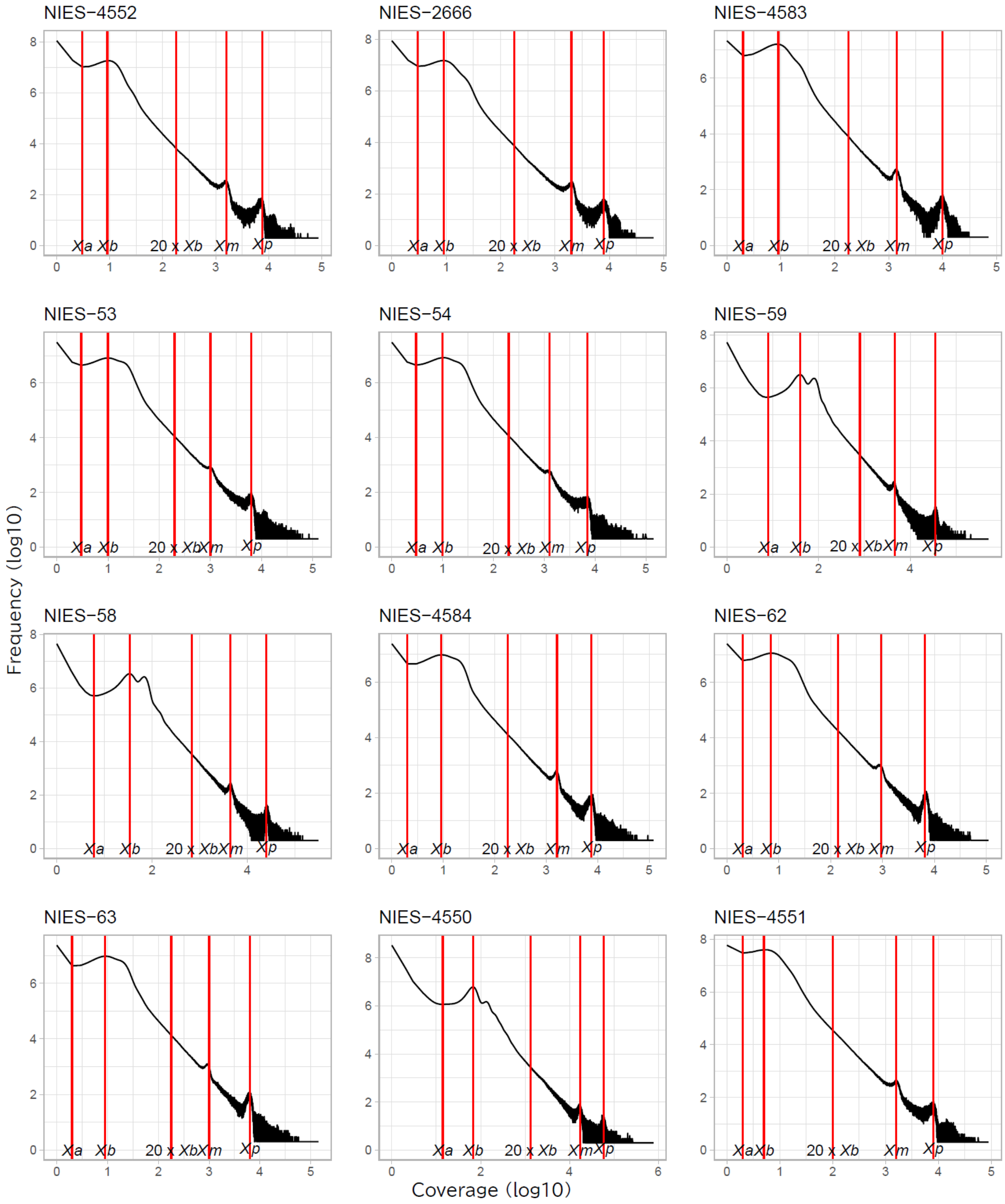
NIES−4552
NIES−2666
NIES−4583
NIES−53
NIES−54
NIES−59
NIES−58
NIES−4584
NIES−62
NIES−63
NIES−4550
NIES−4551
Xa
Xb
20 x Xb
Xm
Xp
Frequency (log10)
Coverage (log10)

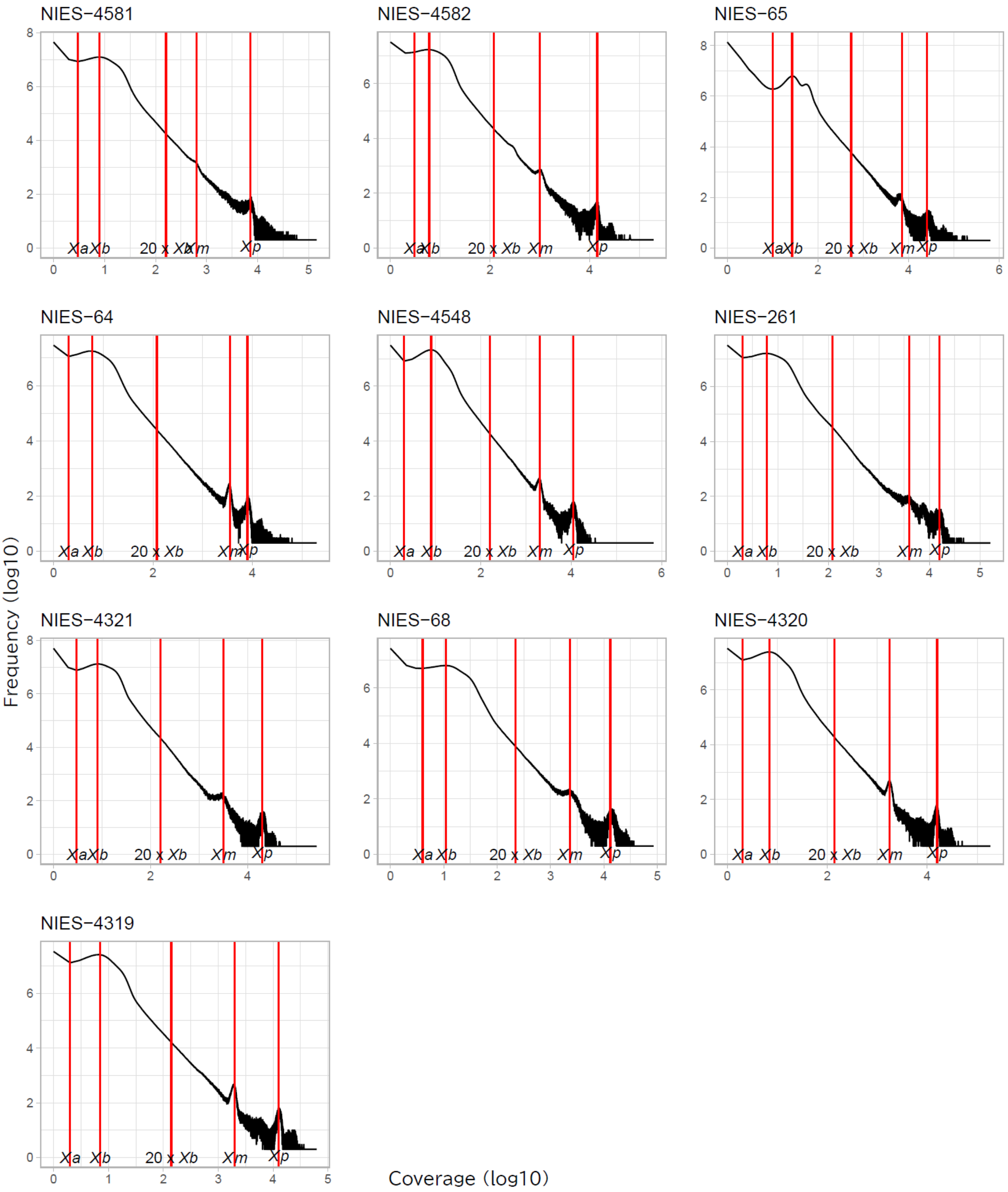


**Fig. S12. Log–log plots of *k*-mer frequency distributions of the 22 strains with *k* = 25.**

Red vertical lines indicate *Xa*, *Xb*, 20 × *Xb*, *Xm*, and *Xp* (from left to right). *Xa*: the first local minimal value. *Xb*: the first peak of the *k*-mer occurrences. *Xm* and *Xp*: peaks of mitochondrial and plastid genomes, respectively.

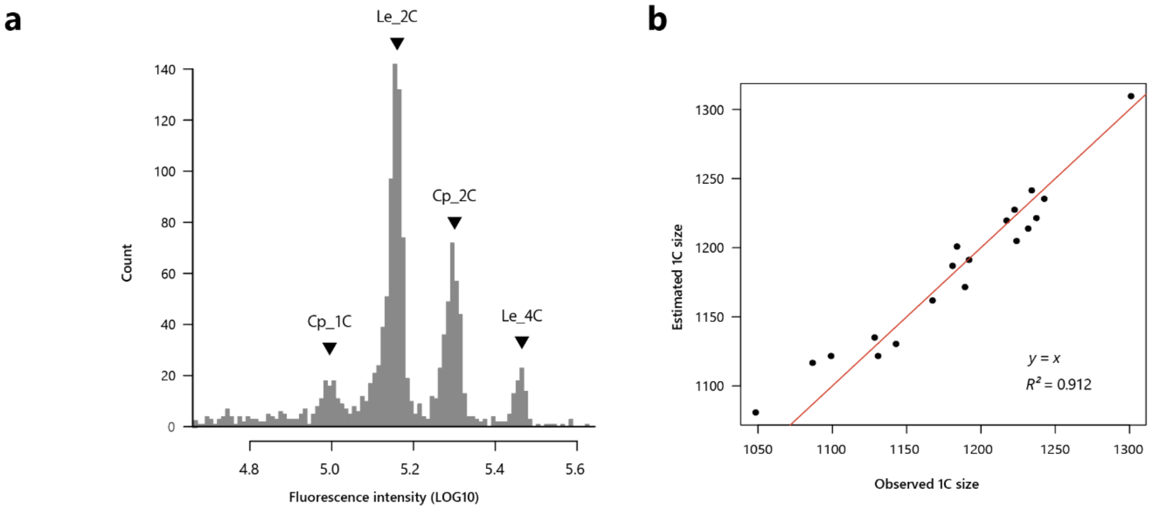


**Fig. S13. Estimation of genome size using flow cytometry.**

(a) Fluorescence intensity of the PI-stained NIES-65 strain and *Lycopersicon esculentum* 'Stupicke' as control. Le_2C and Le_4C indicate the 2C and 4C peaks of *L. esculentum* 'Stupicke', respectively. Cp_1C and Cp_2C indicate the 1C and 2C peaks of NIES-65, respectively. (b) The observed 1C genome size and estimated 1C genome size from the observed 2C genome size.

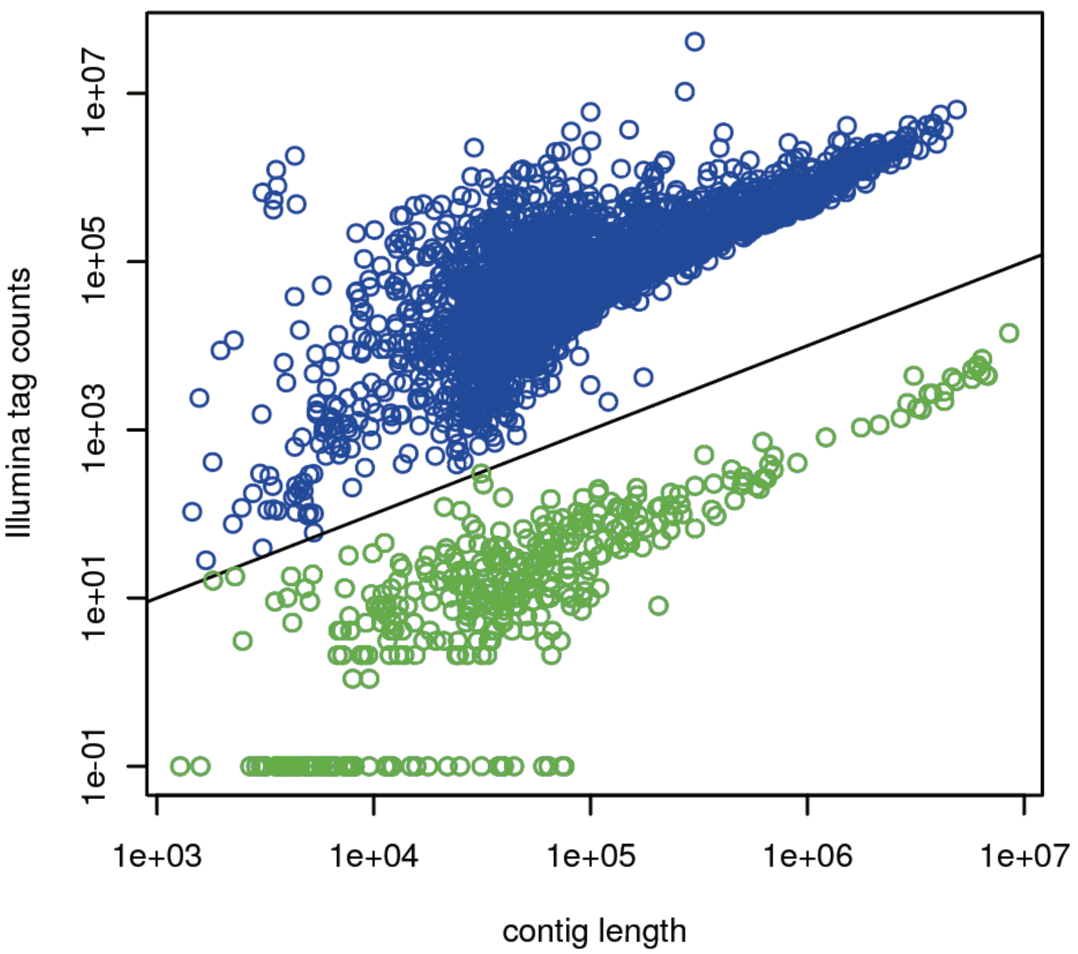


**Fig. S14. The relationship of mapped tag number and the contig length of the genome from the NIES-64 strain.**

Blue and green points are contigs having more and less than 1 tag per 100 bp reference, respectively. Therefore, we retain contigs having more than 1 tag per 100 bp reference as authentic contigs.

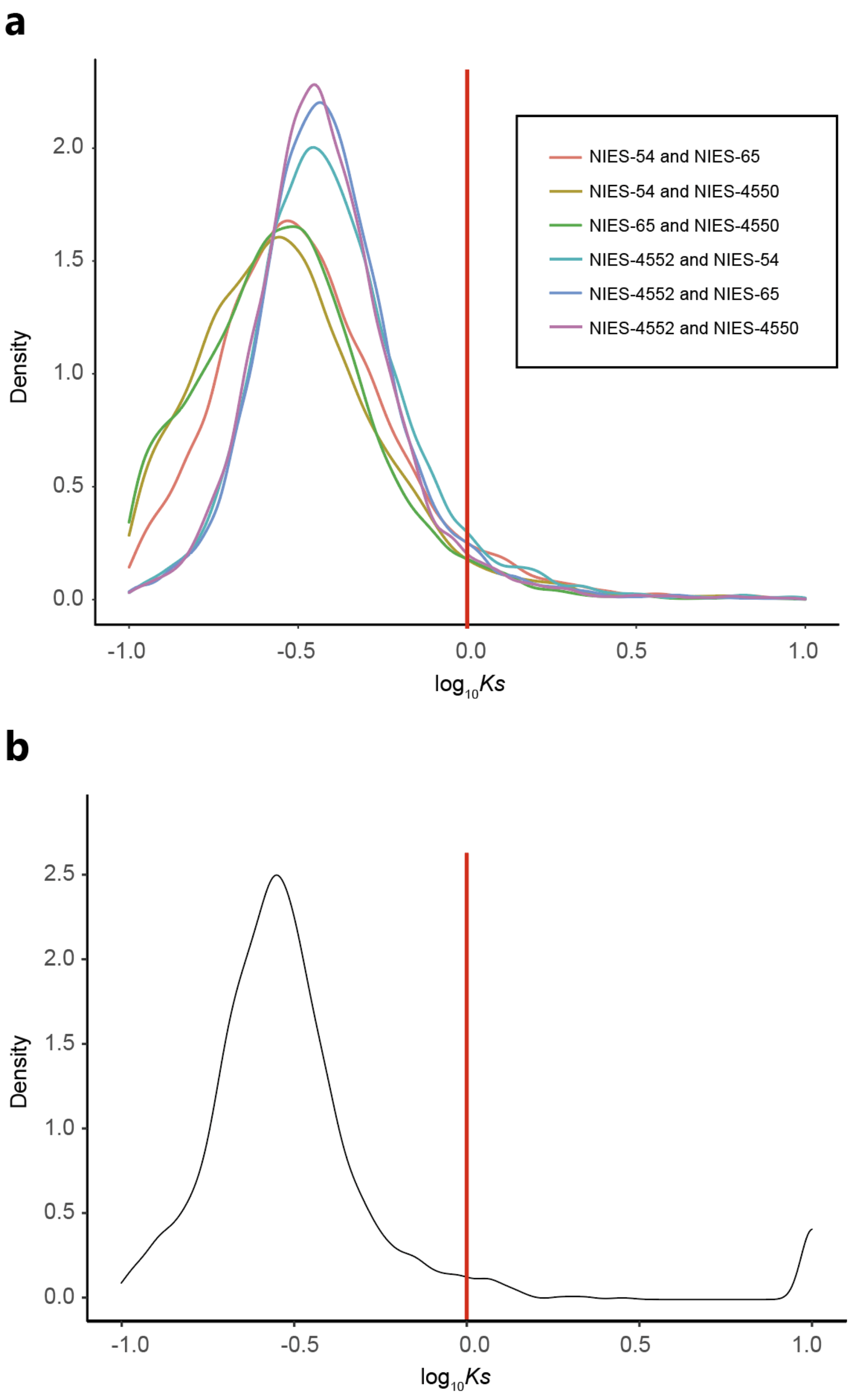


**Fig. S15. *Ks* distribution plot.**

(a) *Ks* distribution plot across all genes from the four strains: NIE-54, NIES-65, NIES-4550 and NIES-4552. (b) The distribution of the maximum *Ks* across all genes within each ortholog group. Red lines indicate *Ks* = 1. We only use OGs with the maximum $K_s$ values < 1 in subsequent analyses.

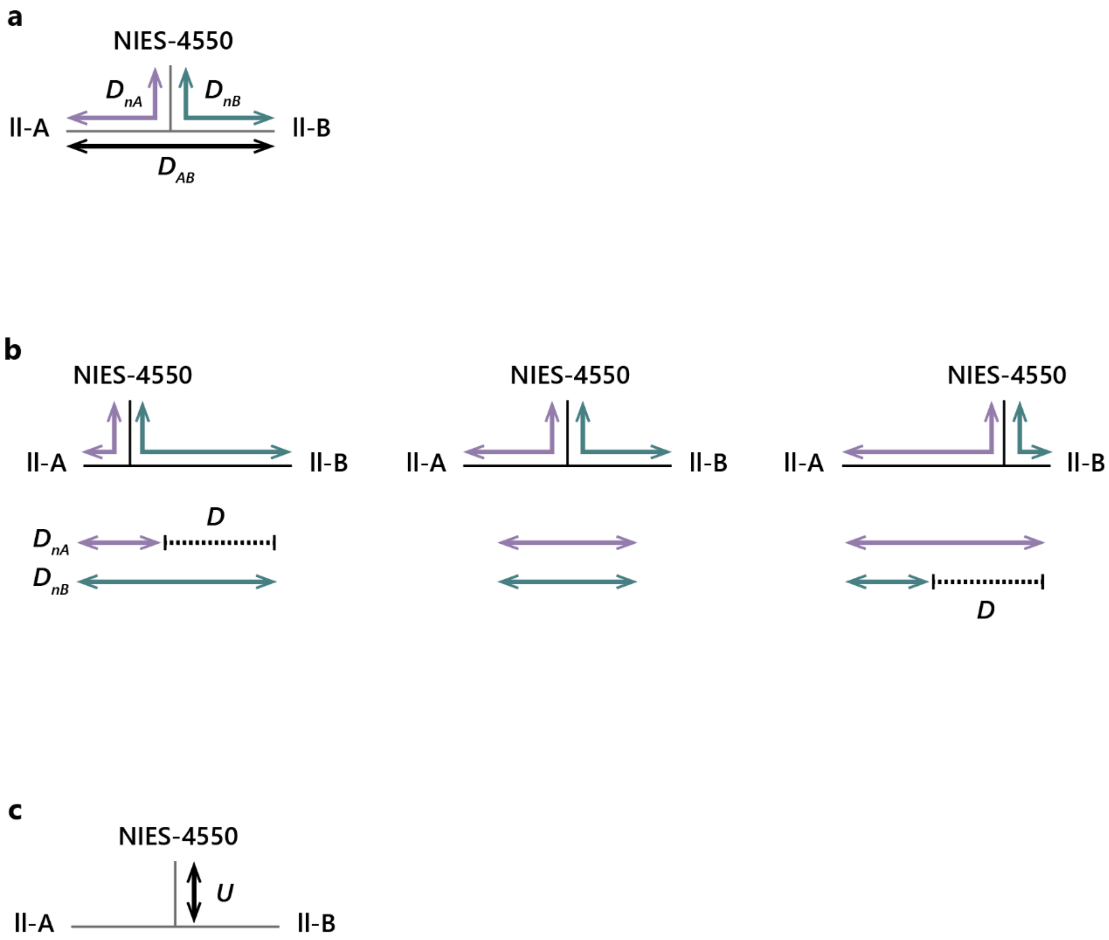


**Fig. S16. Schematic diagrams for the analysis of the origins of the NIES-4550 genome.**

(a) Definitions of $D_{nA}$, $D_{nB}$, and $D_{AB}$. $D_{nA}$ is the average genetic distance of all combinations between the sequences from the NIES-4550 strain and the group II-A. $D_{nB}$ and $D_{AB}$ were also calculated as $D_{nA}$, but for the distance between the NIES-4550 strain and the group II-B and between groups II-A and II-B, respectively. The genetic distance is based on the evolutionary distance calculated by the maximum likelihood method. (b) The definition of $D$ is static, indicating whether the sequence is more closely related to group II-A or II-B. $D$ is the difference between $D_{nA}$ and $D_{nB}$ normalized by $D_{AB}$ to evaluate whether a gene from the NIES-4550 genome is closer to group II-A or II-B. (c) $U$ is the relative number of mutations unique to NIES-4550. If a sequence does not harbor any mutation unique to NIES-4550, $U$ is 0.5.

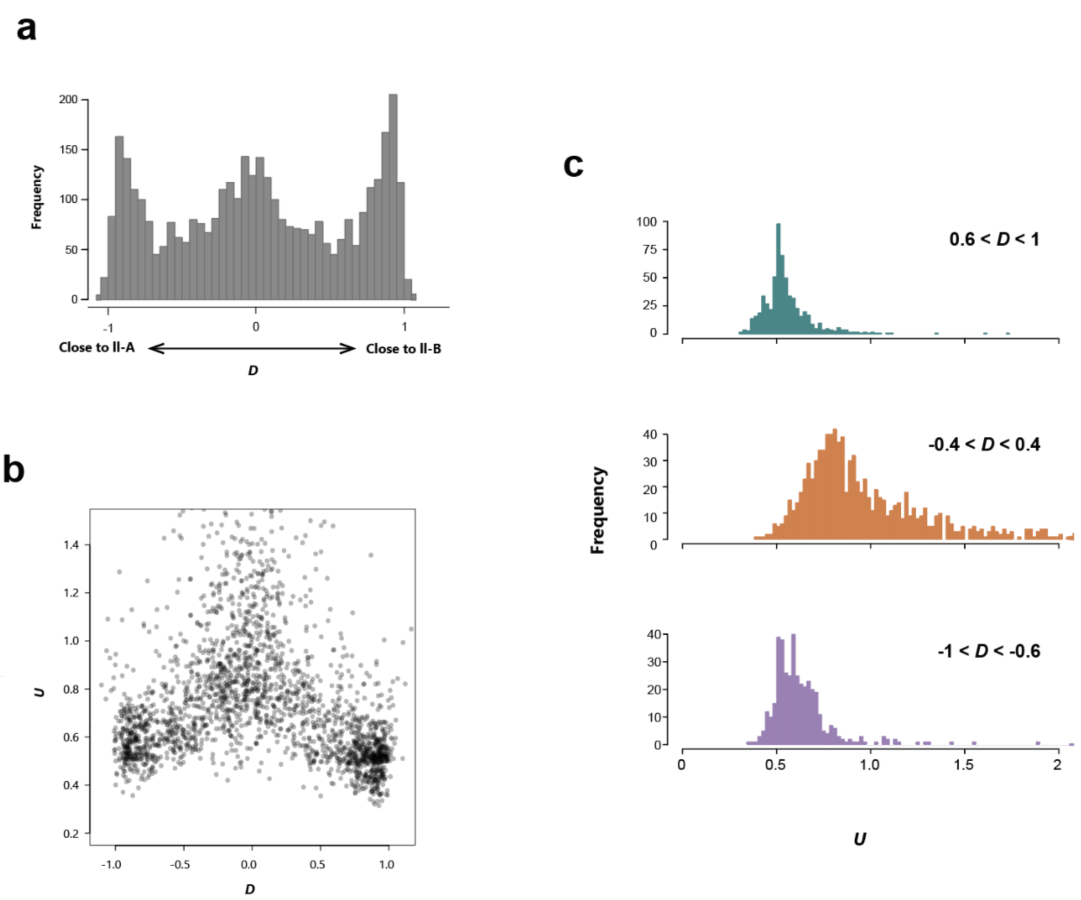


**Fig. S17. Clarifying the origins of the NIES-4550 genome.**

(a) Distribution of *D* of each gene from the NIES-4550 strain. If a gene from NIES-4550 is closely related to the genes from group II-A or II-B, *D* should be smaller or larger than zero, respectively (supplementary fig. S16b left and right). If a sequence from NIES-4550 is highly distant from groups II-A and II-B, *D* should be close to zero. (b) Distribution of *D* and *U* of each gene from NIES-4550. *U* is the relative number of mutations unique to NIES-4550. If a sequence does not harbor any mutation unique to NIES-4550, *U* is 0.5. (c) Distribution of *U* of each gene with different values of *D*.
